# CRISPR-Cas-amplified urine biomarkers for multiplexed and portable cancer diagnostics

**DOI:** 10.1101/2020.06.17.157180

**Authors:** Liangliang Hao, Renee T. Zhao, Chayanon Ngambenjawong, Heather E. Fleming, Sangeeta N. Bhatia

**Affiliations:** Koch Institute for Integrative Cancer Research, Massachusetts Institute of Technology, Cambridge, MA 02139, USA; Institute for Medical Engineering and Science, Massachusetts Institute of Technology, Cambridge, MA 02139, USA; Department of Electrical Engineering and Computer Science, Massachusetts Institute of Technology, Cambridge, MA 02139, USA; Department of Medicine, Brigham and Women’s Hospital and Harvard Medical School, Boston, MA 02115, USA; Broad Institute of Massachusetts Institute of Technology and Harvard, Cambridge, MA 02139, USA; Howard Hughes Medical Institute, Cambridge, MA 02139, USA

**Keywords:** Noninvasive diagnostics, protease activation, CRISPR-Cas, nucleic acid barcodes, urinary fingerprints, point-of-care

## Abstract

Synthetic biomarkers, exogenous probes that generate molecular reporters, represent an emerging paradigm in precision diagnostics with applications across infectious and noncommunicable diseases. In order to achieve their promise, these methods reply on multiplexing strategies to provide tools that are both sensitive and specific. However, the field of synthetic biomarkers has not benefited from molecular strategies such as DNA-barcoding due to the susceptibility of nucleic acids *in vivo*. Herein, we exploit chemically-stabilized DNAs to tag synthetic biomarkers and produce diagnostic signals *via* CRISPR nucleases. Our strategy capitalizes on disease-associated, protease-activated release of nucleic acid barcodes and polymerase-amplification-free, CRISPR-Cas-mediated barcode detection in unprocessed biofluids. In murine cancer models, we show that the DNA-encoded urine biomarkers can noninvasively detect and monitor disease progression, and demonstrate that nuclease amplification can be harnessed to convert the readout to a point-of-care tool. This technique combines specificity with ease of use to offer a new platform to study human disease and guide therapeutic decisions.

## Introduction

Precision diagnostics that reflect the diversity of disease signals are essential for improving therapeutic outcomes. Whereas disease-derived molecules such as proteins or nucleic acids are often limited by either abundance or stability in circulation, rationally-engineered ‘synthetic biomarkers’ leverage the catalytic, functional, and/or microenvironmental nature of enzymes to achieve higher sensitivity and specificity relative to endogenous biomarkers^1–6^. As the development of synthetic biomarkers continues to push molecular diagnostics forward, further progress will necessitate tools capable of achieving highly multiplexed readouts to classify diverse, dynamic disease states.

Molecular strategies for multiplexing abound in analytical chemistry, with DNA-barcoding proving to be transformative in drug screening, gene expression profiling, and cellular function analysis. However, the field of synthetic biomarkers has not yet benefited from DNA multiplexing due to the susceptibility of nucleic acids *in vivo*^7–9^. Even measurements of endogenous, cell-free nucleic acids are hampered by the degradation of sequences that are not protected by the nucleosome in circulation^10–13^. Chemically-modified nucleic acids which have been primarily developed as therapeutics are less resistant to systemic degradation; however, they have not been incorporated into diagnostic platforms because they cannot be amplified with conventional polymerases, nor inexpensively sequenced^14–16^.

To multiplex synthetic biomarkers for use as point-of-care (PoC) precision diagnostics, we exploited chemically-stabilized DNA for molecular barcoding and employed CRISPR-Cas nucleases that are insensitive to chemical modifications to read DNA barcodes in a sequence-specific manner^17^. By integrating these enabling technologies, we engineered a panel of high-throughput, programmable *in vivo* sensors for noninvasive disease detection and monitoring in urine (Fig. 1). These rationally-designed, DNA-encoded synthetic urine biomarkers (DNA-SUBs) 1) target sites of disease, 2) sense dysregulated proteolysis^18,19^, and 3) emit nucleic acid barcodes that survive circulation and concentration by the kidney to 4) activate single-stranded DNase activity of CRISPR-Cas12a *via* sequence complementarity to a CRISPR guide RNA. Shed by pathological proteolytic activities, chemically-stabilized DNA barcodes were readily scanned in unprocessed urine using fluorescence kinetics or on paper strips upon Cas12a activation. To demonstrate the modularity of this approach, we assembled DNA-SUBs on both biologic (nanobody) or synthetic (polymer) scaffolds to demonstrate their use in noninvasive monitoring of two types of invasive cancer.

**Figure 1.**
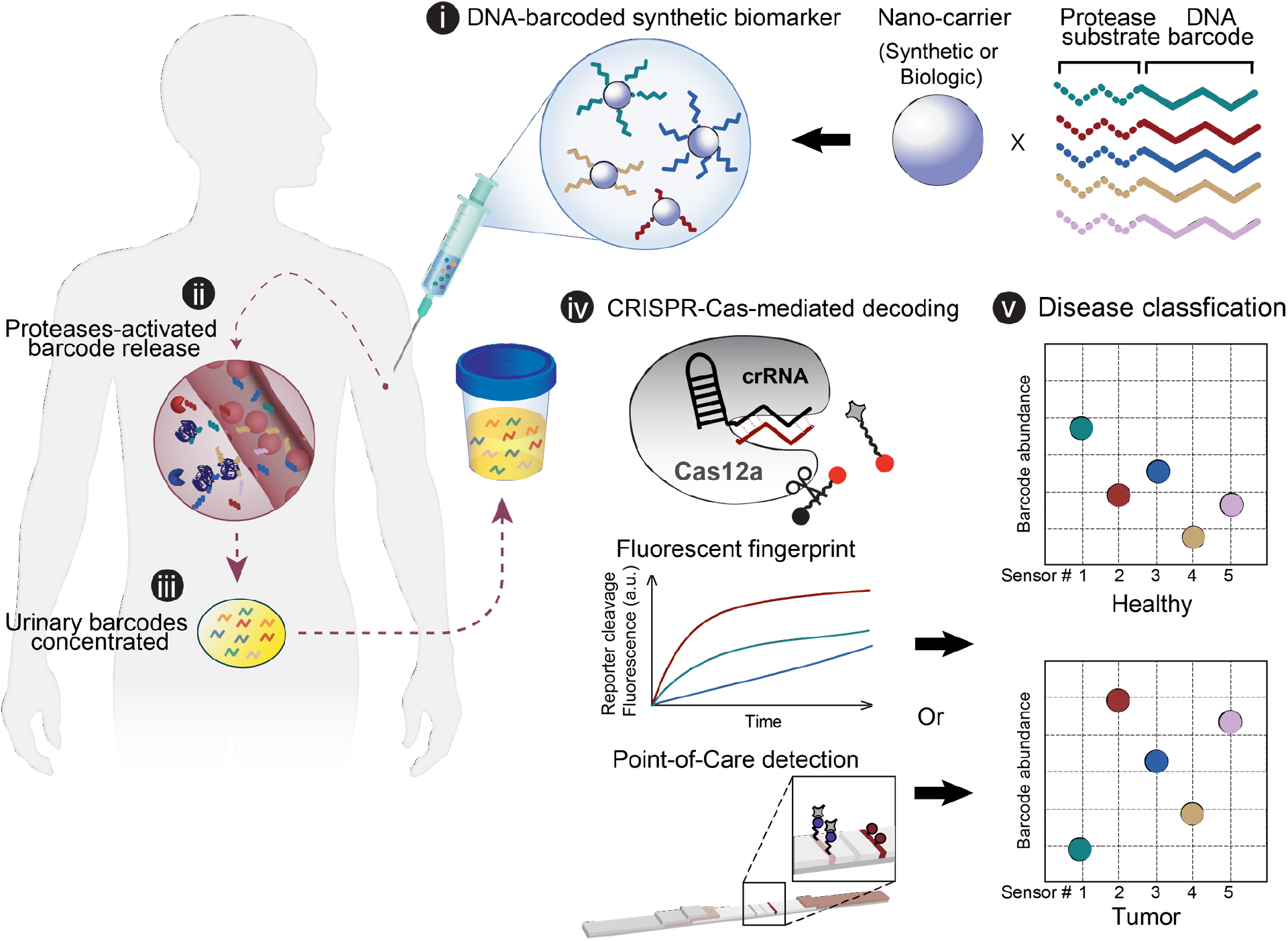
Engineering DNA-encoded synthetic urine biomarkers with CRISPR-Cas-mediated disease detection. DNA-encoded synthetic urine biomarkers are comprised of a nano-carrier (synthetic or biologic) functionalized with protease-activated short peptides barcoded with oligonucleotides (i). After *in vivo* administration, activation of DNA-encoded sensors by disease-specific protease activity triggers release of synthetic DNA barcodes (ii) that are size-specifically concentrated in the urine for disease monitoring (iii). DNA barcodes in the urine activate programmable CRISPR-Cas nucleases to release the multiplexed reporter signals that are fluorescent, or detected on paper (iv), enabling *in situ* disease classification at the point-of-care via the patterns of local proteolytic activities in the disease microenvironment (v).

## Results

### Chemically-modified DNA enables CRISPR-Cas12a-mediated urinary readout for *in vivo* sensing

CRISPR-Cas nucleases show great promise in DNA and RNA diagnostic applications^17,20–25^, though they have yet to be applied to *in vivo* disease sensing due to the unspecific degradation of unmodified nucleic acids in circulation. To prevent the susceptibility and immunostimulatory activity of unmodified nucleic acids *in vivo*, we sought to investigate the tolerability of Cas proteins to chemically-stabilized DNA molecules. We introduced native and FDA-approved phosphorothioate-modified DNA molecules to the enzyme CRISPR-Cas12a^26–28^. The Cas12a from *Lachnospiraceae bacterium* ND2006 (*Lba*Cas12a) assembled with guide CRISPR RNA sequences (crRNAs) recognizes 1) a T nucleotide-rich protospacer-adjacent motif (PAM) to target double-stranded DNA (dsDNA), or 2) single-stranded DNA (ssDNA) through sequence complementarity in a PAM-independent manner, and unleashes a robust, nonspecific ssDNA trans-cleavage activity that can be monitored using a fluorophore (F)- quencher (Q)- labeled reporter^17^ (Fig. 2A). In addition to native dsDNA or ssDNA, *Lba*Cas12a was activated by phosphorothioated ssDNA at a relatively slower speed (Fig. 2B). When intravenously administered into a murine model (Balb/c mouse), native ssDNA in urine collected from injected animals could not activate *Lba*Cas12a assembled with the corresponding crRNA due to the unspecific DNase activities in circulation (Fig. 2C, Fig. S1A). In contrast, different lengths of phosphorothioate-modified ssDNAs in solution or unprocessed urine from injected animals triggered the trans-cleavage activity of *Lba*Cas12a (Fig. 2C). Notably, the 20-mer crRNA-complementary ssDNA optimized kidney filtration into urine, producing the highest reporter cleavage activity, whereas the 24-mer ssDNA containing the PAM sequence produced the highest cleavage signal *in vitro* (Fig. 2C, Table S1, S2). Furthermore, we validated multiple crRNA-modified ssDNA activator pairs with orthogonality between different sequences, allowing for parallel readout in multiple well assays (Fig. 2D, Fig. S1B-H). The *Lba*Cas12a was activated once it encountered its programmed DNA target in unprocessed urine and cleaved a FAM and biotin dual-tagged ssDNA reporter that rapidly appeared on a lateral flow paper strip (Fig. 2E). The presence of the ‘sample band’ at the front of the strip indicated that the cleaved reporters were produced upon the activation of *Lba*Cas12a by the urinary DNA activator. In addition to the binary DNA activator detection, quantification of the intensity of the sample bands on paper strips allowed assessment of enzymatic kinetics (Fig. 2F). By adjusting the concentrations of assay components, the working linear ranges fell within sub-nanomolar DNA activator concentrations for both fluorescent and paper-based readouts (Fig. S2, S3).

**Figure 2.**
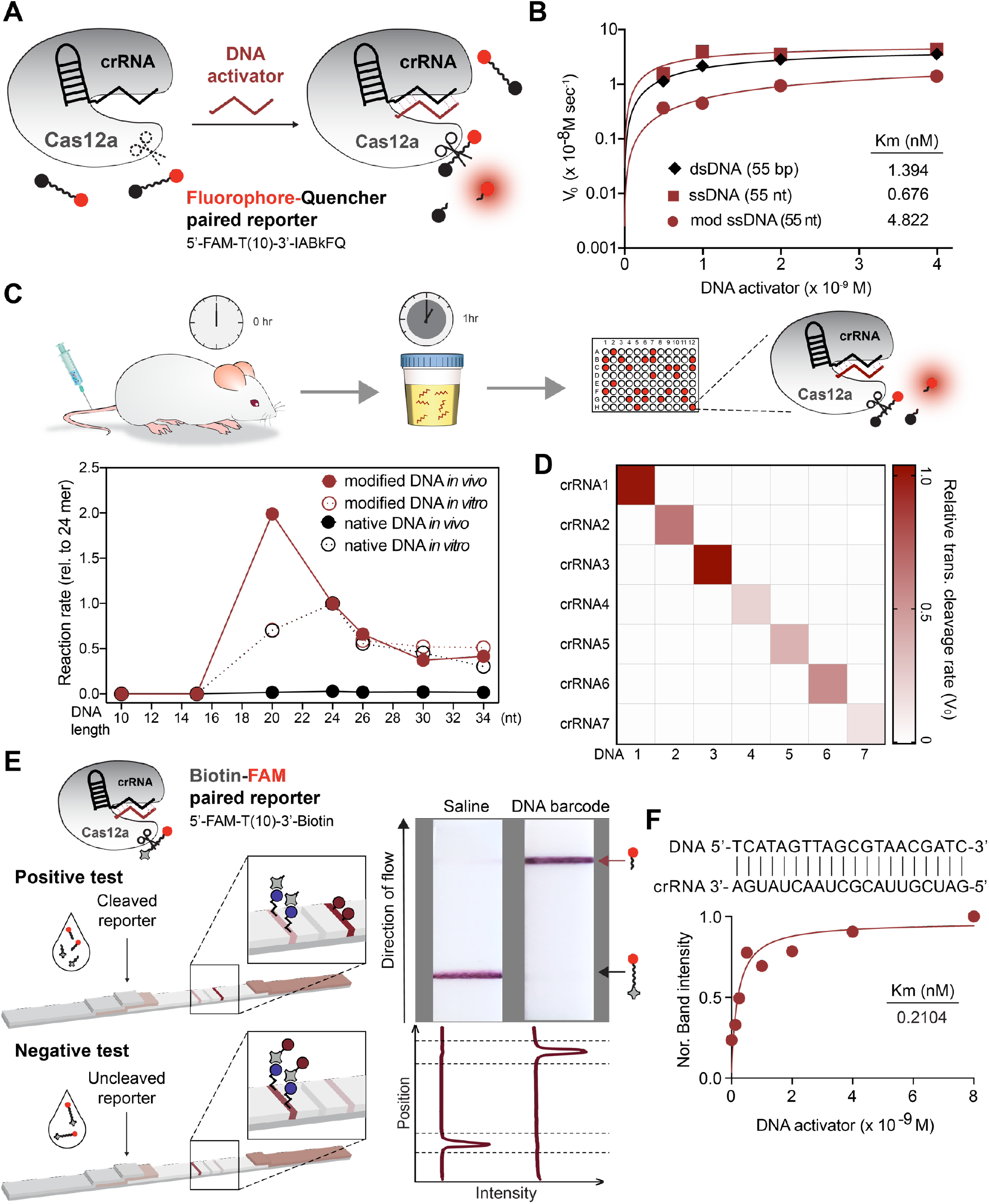
Chemically-modified DNA enables CRISPR-Cas-mediated urinary readout for *in vivo* sensing. (A) DNA fragment activates nonspecific single-stranded DNase cleavage upon binding to crRNA on Cas12a. Such activity can be tracked by the dequenching of fluorescence from a fluorophore (F, 5’-FAM)-quencher (Q, 3’-IABkFQ)-paired oligonucleotide. (B) Representative Michaelis-Menten plot of LbaCas12a-catalyzed ssDNA trans cleavage using a native dsDNA, ssDNA, or fully phosphorothioate-modified ssDNA activator. The initial reaction velocity (V0) is determined from the slope of the curve at the beginning of a reaction. (C) Schematic showing urine testing in a mouse model and the study time course (1 h). Plot underneath shows length optimization of ssDNA activator *in vitro* and *in vivo*, by quantifying the trans-cleavage rate of Cas12a upon activation of native or modified ssDNA in solution (4 nM) or mouse urine (1 nmol per injection). The trans-cleavage rates in each condition in the format of initial reaction velocity were normalized to that of a 24-mer (See also Table S2). (D) Heatmap of trans-cleavage rates of different modified ssDNA activator-crRNA pairs. Assays were performed with urine samples collected from mice injected with 1 nmol of modified ssDNA activator after 1 h of *i.v.* administration. (E) Schematic showing set up of paper-based lateral flow assay (LFA) (left). When Cas12a is activated by the DNA activator in mouse urine, it cleaves the fluorescein (FAM)-biotin-paired oligonucleotide reporter and frees the FAM molecule that can be detected on the ‘sample band’. Uncleaved reporters are trapped on the ‘control band’ via binding of biotin to streptavidin. Different bands are visible on paper strips (right). Band intensities were quantified using ImageJ, and each curve was aligned below the corresponding paper strip. The top peak of the curve shows the freed FAM molecule in cleaved reporter samples, and the bottom peak shows the presence of the uncleaved FAM-biotin reporter. (F) Michaelis-Menten plot of LbaCas12a-catalyzed ssDNA trans-cleavage upon a representative DNA-crRNA pairing (complementary sequences are shown) on paper. Data were plotted with the quantified band intensity of cleaved reporter on paper strips (See also Figure S1-3).

### Tumor targeting, singleplex DNA-encoded synthetic urine biomarker specifically detects tumor-bearing animals

To develop DNA-encoded synthetic biomarkers, we leveraged deregulated proteolytic activities in the disease microenvironment to cleave and release the phosphorothioated DNA barcodes that were size-specifically concentrated in the urine to produce a noninvasive readout of the target disease. We first evaluated a singleplex synthetic biomarker *in vivo* in a human prostate cancer (PCa) xenograft model^29^. To maximize the on-target protease cleavage, we engineered the DNA-SUB on a biological scaffold that enables tumor-targeting abilities. To utilize the robust stability and tissue affinity of single domain antibody fragments (nanobodies), we constructed DNA-encoded, protease-activatable nanobodies by inserting a peptide substrate sequence with an unpaired cysteine for one-step site-specific labeling of cargos *via* a thio-ether bond^30–33^ (Fig. 3A, Table S3, Fig. S4A, B). The peptide substrate specifically responded to the PCa-associated protease PLAU^29^ (Fig. S7E). To prevent possible misfolding caused by the internal disulfide bond present in the nanobody, the peptide substrate with cysteine was spaced from the nanobody scaffold by a rigid linker (Table S3). For *in vivo* validation, we tested a PLAU-activated, cMET-targeting nanobody in the cMET- and PLAU-expressing PC-3 cell-derived tumor model. In the subcutaneous PC-3 tumors, the cMET nanobody mediated active tumor trafficking upon systemic administration (Fig. 3B, Fig. S4C, D). PLAU-activated nanobodies covalently conjugated with the 20-mer DNA barcode were efficiently separated *via* size-exclusion chromatography. The DNA-barcoded, PLAU-activated cMET nanobody (cMET-Nb-DNA) exhibited enhanced tumor accumulation compared with the DNA-barcoded, PLAU-activated non-targeting GFP nanobody (GFP-Nb-DNA)^34^ (Fig. 3C). We systemically administered cMET-Nb-DNA to tumor-bearing and healthy control mice and quantified urinary DNA barcodes that were freed from the nanobody scaffold 1 h after injection. Urine samples were incubated with *Lba*Cas12a-coupled with the complementary crRNA, and the trans-cleavage activity triggered by the DNA barcode was analyzed by tracking the kinetics of cleavage of a fluorescence-quencher labeled poly(T) reporter (Fig. 3D). Administration of cMET-Nb-DNA significantly increased the trans-cleavage rate of *Lba*Cas12a activated by urine samples collected from tumor-bearing mice, relative to that of the healthy controls (Fig. 3C). To translate the fluorescent readout into a PoC detection tool, we incubated *Lba*Cas12a activated by mouse urine samples with the FAM-poly(T)-biotin reporter and ran the cleavage products on lateral flow paper strips. An enhanced sample band appeared in samples collected from tumor-bearing mice injected with cMET-Nb-DNA (Fig. 3D). The high sensitivity and specificity of the sensor to track disease was reflected in a ROC curve (AUC= 0.89). In contrast, urine samples collected from tumor-bearing mice injected with GFP-Nb-DNA activated *Lba*Cas12a at a rate that was almost identical to the samples from healthy controls, indicating that the release of the DNA barcodes was triggered on-target (Fig. 3D, E).

**Figure 3.**
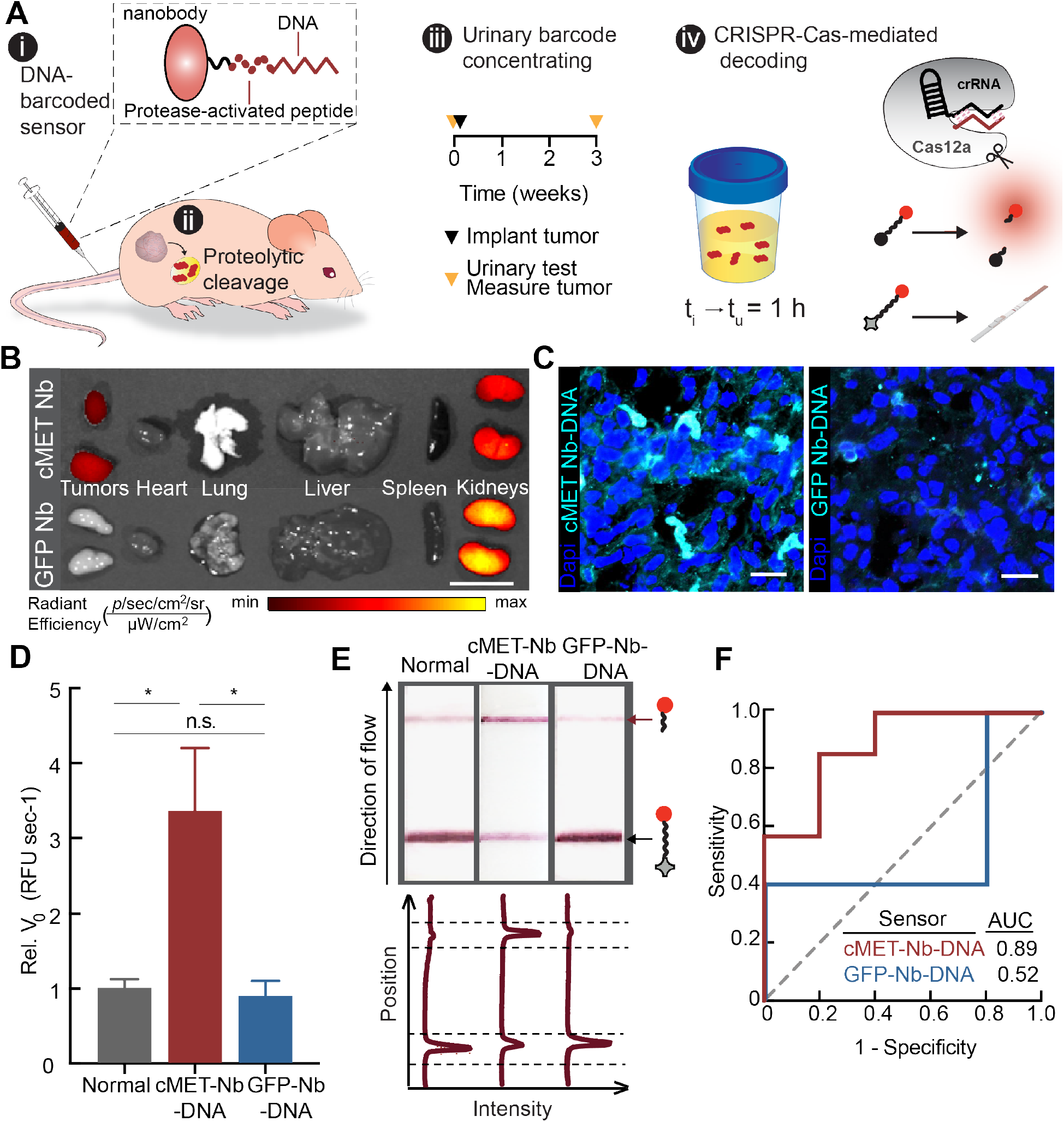
Tumor-targeting, singleplex DNA-encoded synthetic urine biomarker for detection of prostate cancer. (**A**) Schematic showing urine testing in a human prostate cancer xenograft model: generation of the DNA-encoded protease-activatable nanobody with an unpaired cysteine and 1-step conjugation of ssDNA activator (i). Activation of diagnostic by disease-specific protease activity triggers (ii) release of ssDNA activator into urine for disease detection (iii). Study time course of urine testing and detection of the ssDNA activator with Cas12a trans-cleavage assay using fluorescent or paper readout (iv). *t*, time at given testing step; *t*_*i*_, time of sensor injection; *t*_*u*_, time of urine collection. (**B**) IVIS image shows biodistribution of cMET targeting nanobody and on-targeting GFP nanobody when injected intravenously in nude mice bearing PC-3 xenografts. Scale bar = 1 cm. (**C**) Immunofluorescent staining of Cy7-labeled DNA-encoded cMET nanobody and DNA-encoded, non-targeting GFP nanobody on sections of PC-3 tumors. Scale bar = 20 μm. (**D**) Unprocessed urine samples collected from tumor-bearing mice injected with DNA-encoded cMET nanobody or DNA-encoded GFP nanobody, and healthy control mice injected with DNA-encoded cMET nanobody were applied in the Cas12a trans-cleavage assay. Initial reaction velocity (V_0_) of the Cas12a trans-cleavage assays were calculated and normalized to that of healthy control mice (n=5 or 7 mice per group; ± SEM; unpaired *t*-test with Welch’s correction, \**P*<0.05). (**E**) Paper-based LFA of Cas12a activated by urine samples collected from tumor-bearing or healthy control mice in (C). Band intensities were quantified using ImageJ and each curve was aligned below the corresponding paper strip. The top peak of the curve shows the presence of the cleaved reporter and the bottom peak shows the presence of the uncleaved reporter. (**F**) ROC curves characterize the predictive power of a biomarker by returning the area under the curve (AUC) as a metric, with a baseline AUC of 0.5 representing a random biomarker classifier. AUC comparison between DNA-encoded cMET nanobody or DNA-encoded GFP nanobody injected tumor cohort against normal cohort in (C). Dashed line represents an AUC of 0.5, and a perfect AUC is 1.0.

### DNA-encoded multiplex synthetic urine biomarkers longitudinally monitor disease progression in a portable manner

It is increasingly appreciated that analysis of multiple cancer hallmarks may optimize diagnostic sensitivity and specificity in heterogenous diseases. Whereas active targeting is limited to diseases that express specific ligands, multiplexing of an untargeted scaffold has the potential to be more generalizable. Therefore, we constructed a multiplexed panel of DNA-SUBs on a polymer-based scaffold and administered them as a single pool to mice (Fig. 4A). Each DNA-SUB was comprised of a 20-mer phosphorothioated DNA-tagged, protease-activated peptide (PAP) covalently conjugated to a synthetic polymer (8-arm polyethylene glycol, 40 kDa) (Fig. 4A, Fig. S5 & Table S3). To monitor nanosensor trafficking to tissue contact, we established a syngeneic mouse model by intravenously injecting a metastatic murine colorectal cell line (MC26-LucF) into immunocompetent Balb/c mice^35^ (Fig. 4A, Fig. S6). We first identified a panel of CRC-specific proteases through transcriptomic analysis and found multiple proteases expressed in tumors at >1.5-fold levels over normal samples, including members of the matrix metalloproteinase (MMPs), aspartic, and serine protease families (i.e. cathepsins, kallikrein-related peptidases) (Fig. S7A). From a matrisome proteomic analysis, we confirmed proteases present in primary CRCs and their distant metastases (e.g. MMP-7, -9, Cathepsin D, PLAU)^36,37^ (Fig. S7C). We confirmed that these identified proteases were overexpressed in tumor-bearing lung tissue of the MC26 transplantation model compared to normal lung tissue (Fig. S7B, D). To identify peptide substrates specific to the selected proteases, we screened 16 peptide sequences against purified recombinant proteases and identified the top five substrates using a fluorogenic activity assay^29,38^ (Fig. S7E). These protease-activated peptides (PAP7, PAP9, PAP11, PAP13, PAP15) broadly cover metallo, serine, and aspartic protease activities (Fig. S7E), and were specifically cleaved by tumor tissue homogenates *ex vivo* with high predicted disease classification power, and thus were incorporated into the panel of DNA-SUBs for *in vivo* validation (Fig. 4B, Fig. S8).

**Figure 4.**
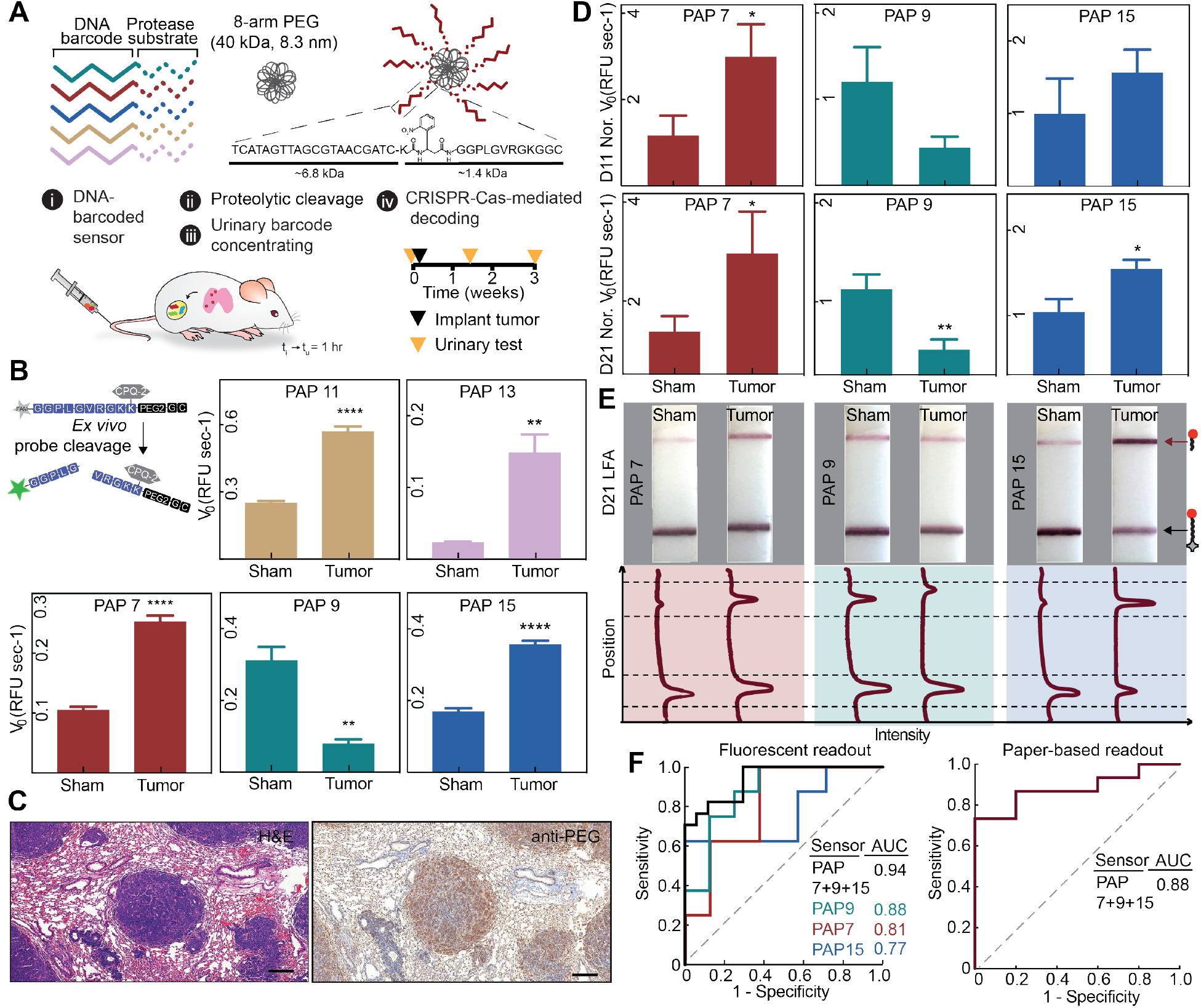
DNA-encoded multiplex synthetic urine biomarkers for portable monitoring of invasive CRC. (**A**) Scheme of the work flow for longitudinal disease monitoring with the multiplexed DNA-encoded synthetic urine biomarkers. (**B**) Forster resonance energy transfer (FRET)-based peptide assay to identify the real-time cleavage of peptide substrates by invasive CRC tissue homogenates collected 21 days after tumor inoculation. Peptide cleavage kinetics were monitored and cleavage rates were plotted (n=5 mice per group; ± SEM; unpaired *t*-test with Welch’s correction, \*\**P*<0.01, \*\*\*\**P*<0.0001). (**C**) Histological staining of lung sections of Balb/c mice bearing CRC lung tumor nodules 21 days after tumor inoculation (left). Mice were injected with 5-plex DNA-SUBs with the PEG core and immunohistochemistry of the same tissue stained with anti-PEG antibody (right). Scale bar = 200 μm. (**D**) Pooled DNA-SUBs were administered to Balb/c mice bearing CRC lung tumor nodules (tumor) and saline-injected control animals (sham) at day 11 or 21 after tumor initiation. All urine samples were collected at 1 h after sensor administration. Cas12a trans-cleavage assays were performed against each DNA-barcode with the fluorophore-quencher paired reporter. Initial reaction velocity (V_0_) of the Cas12a trans-cleavage assays were calculated and normalized to that of saline injected control animals (n=8 or 10 mice per group; ± SEM; unpaired *t*-test with Welch’s correction, \**P*<0.05, \*\**P*<0.01). The initial reaction velocity (V_0_) refers to the slope of the curve at the beginning of a reaction. (**E**) Representative paper strips of the paper-based LFA of Cas12a activated by mouse urine samples collected in (D). Band intensities were quantified using ImageJ. The top peak of the curve shows the freed FAM molecule in cleaved reporter and the bottom peak shows the presence of the uncleaved FAM-biotin reporter. (**F**) Left, ROC curve analysis indicates predictive ability of single or combined DNA-SUBs with fluorescent readout in (D). Right, ROC curve shows the predictive ability of paper-based urinary readout in (E). ROC analysis utilized ratio of quantified cleaved reporter band intensity over its corresponding control band intensity. Dashed line represents an AUC of 0.5, and a perfect AUC is 1.0.

We first showed that a DNA-barcoded, MMP-responsive SUB (DNA-PAP7-SUB) accumulated in the CRC lung tumor nodules following intravenous injection (Fig. 4C, Fig. S9B). We then tested the entire 5-plex of DNA-barcoded SUBs *in vivo*, with an emphasis on identifying reporters that differentiated mice bearing lung tumor nodules from the healthy control animals. We systemically administered the multiplexed DNA-SUBs to the two mouse cohorts over the course of tumor development, and quantified urinary DNA barcodes that were freed from the nanosensors one hour after injection. Urine samples were incubated with *Lba*Cas12a-coupled with five different complementary crRNAs in multiple wells, and the trans-cleavage activity triggered by each DNA barcode was analyzed by tracking the cleavage kinetics of a fluorescence-quencher labeled reporter. We found that the MMP-responsive sensor (DNA-PAP7-SUB) from this multiplexed panel succeeded in distinguishing tumor-bearing mice from healthy mice only 11 days after tumor inoculation when the tumor nodules were 1-2 mm. Some sensor (DNA-PAP9-SUB, DNA-PAP15-SUB) differences were amplified over time (Fig. 4D, Fig. S6A, Fig. S9C), and these sensors (PAP9 and PAP15) corresponded to peptides cleaved by serine and metalloproteases *in vitro*, and also produced distinct cleavage patterns when incubated with homogenates from either tumor-bearing or healthy lung tissues (Fig. 4B). Based on the ROC curve analysis, the sum of the metallo (PAP7, PAP15) and serine (PAP9) protease substrate signals significantly increased the classification power of the DNA-SUBs (combined sensors PAP7/9/15 AUC=0.94; PAP7 AUC= 0.81; PAP9 AUC=0.88, PAP15 AUC=0.77; Fig. 4F). We then combined the 5-plex sensor panel with lateral flow detection for a visual readout that could enable PoC diagnostics. Using the same urine samples assayed in the aforementioned Cas12a kinetic cleavage reactions (Fig. 4D), we designed the lateral flow assay to read the cleavage of the FAM-poly(T)-biotin reporter at the optimized end timepoint. After the activation of *Lba*Cas12a by incubating the enzyme with a specific crRNA and its complementary DNA barcodes in urine, the enzyme complex and FAM-poly(T)-biotin reporter were mixed and added onto an assigned location in a 96-well plate. A series of lateral flow strips were loaded onto the plates and the multiple-pot test paper results appeared in 5 min at room temperature (Fig. 4E). Consistent with the results in the fluorescent readout, the test paper ‘fingerprints’ revealed distinctions in the intensity of sample bands resulting from Cas12a activation of tumor-bearing mice and healthy mice (Fig. 4E, Fig. S9D). Notably, quantification of the sample band intensities exhibited disease classification power with multiple sensors (Fig. 4F), enabling a platform that is amenable to clinical translation due to its well-understood chemical composition and use of DNA multiplexing to overcome relatively low tumor accumulation, relative to ligand-targeted scaffolds.

## Discussion

In this work, we present an integrated synthetic biomarker platform that enables *in vivo* sensing of endogenous proteolytic activities in the disease microenvironment and releases DNA fingerprints that are detectable in the urine. These size-specifically concentrated (< 10 kDa), chemically-stabilized DNA molecules triggered a second signal amplification step through activation of a CRISPR-Cas nuclease in a PAM-independent manner. The pathologically-released synthetic urinary DNAs fulfill several required characteristics of an ideal biomarker: high signal-to-noise ratio, stability in circulation until detection, easy accessibility from host biofluids, and high sensitivity and specificity in disease discrimination. Despite the lower melting temperature of duplexes, the phosphorothioate internucleotide linkages endowed the DNA barcodes with nuclease resistance that was essential for *in vivo* sensing. crRNA complementarity was critical for catalytic activation of *Lba*Cas12a by a DNA activator in solution or injection *in vivo*. Additionally, the molecular weight of the DNA activators played an important role *in vivo* since they underwent characteristic single-exponential concentration decay after intravenous injection followed by size-dependent renal filtration from the blood (< 10 kDa, peaking at 1 hour after delivery)^39^. Therefore, the ~7 kDa 20-mer DNA activator that effectively concentrated in urine outperformed longer PAM-containing sequences in trans-ssDNA cutting. In localized tumors, naturally-occurring ctDNAs are often present at a low abundance and require extensive enrichment steps prior to sequencing. In CRISPR-based nucleic acid detection approaches, the inputs are samples amplified by PCR or Recombinase Polymerase Amplification (RPA). At a tolerable diagnostic dose (<0.3 mg/kg), the synthetic DNA barcodes are readily accessible from compositionally simple, unprocessed urine. The DNA detection sensitivity can be further improved by optimizing the ssDNA trans-cleavage activity of *Lba*Cas12a through enhanced base pairing between the DNA activator and modified crRNA.

In the synthetic urinary biomarker platform, release of DNA barcodes is triggered by proteases known to respond during tumorigenesis. We identified proteases that were differentially expressed in two types of invasive cancer and selected peptide substrates with broad protease susceptibility. To increase the specificity of *in vivo* sensing, we genetically encoded protease-activated sequences fused to a nanobody to selectively bind an overexpressed growth factor receptor in prostate cancer. In addition to the diagnostic demonstration in flank xenograft cell line models, we showed disease classification by multiplexing DNA in a syngeneic model to embrace proteolytic contributions from immune cells. We observed the expected preferential cleavage of metalloprotease-responsive nanosensors in CRC-bearing mice and the activation of a serine protease-sensitive substrate (PAP9) that was reported in the lung. This pattern could point to an imbalance of serine protease/protease inhibitors in the pulmonary tumor microenvironment^40–42^. Due to the differences in proteolytic networks present in mouse models and human diseases, additional work to be completed before clinical translation may include an evaluation of the sensor performance in the context of additional animal models with comorbidities, and also in human samples.

Multiplexing also contributed to enhanced the sensitivity of activity-based nanosensors in previous work by our group. Urinary detection of cancers in mice *via* profiling protease activities by mass spectrometry showed improved sensitivity relative to existing and emerging blood-based diagnostics^1,38^. Owing to the inherent programmability of Cas proteins, Cas12a exhibited collateral activity upon multiple 20-mer barcodes tested, which can be extensible to hundreds of orthogonal codes, providing a level of multiplexing that is challenging to attain with commercial isobaric tags for mass-spectrometry^1^. We thus envision that the DNA-encoded synthetic biomarkers may be used to sense and report the entire activity landscape of proteases in order to differentiate between health and disease, in combination with spatial transcriptomic and multiplexed immunohistological techniques.

Despite the advantages of multiplexing with mass encoding, one challenge of mass spectrometry is that it requires rigorous assessment of instrumentation and data interpretation. Here, through DNA-barcoding, our platform expanded the application of cost-effective CRISPR-based detection from viral or endogenous nucleic acids to synthetic biomarkers^17,20,23,25,43^. This study represents a step towards cancer diagnostics through a lateral flow readout of the synthetic DNA in urine, allowing for rapid, portable detection on paper strips similar to a home pregnancy test. Further refinement of nanosensor formulation and dosing could enhance the performance of SUBs and improve disease management and outcomes for patients. In addition to a systemic injection that is widely available in clinical settings, alternative delivery routes of synthetic biomarkers include oral delivery, topical delivery, or wearable devices, and would allow sensor administration without trained phlebotomists. We therefore anticipate greater accessibility for patients to constantly self-monitor their own disease progression to enable early detection and access to effective treatment.

The specialized substrate recognition and signal amplification of enzymes allows the DNA-encoded SUBs to be a fully modular system with both a designable input *via* a protease-responsive linker and a programmable output *via* CRISPR-Cas activity. We improved the on-target rate of diagnostics by using functional biologics that can be extended to scaffolds with therapeutic efficacy, providing an opportunity to combine therapy and diagnostics to monitor treatment responses in real-time^44^. Future expansion to additional enzymatic families will provide a new noninvasive way to elucidate multi-enzymatic networks at a systemic level^45^. Through tailored target specificities, these programmable sensors may monitor other infectious and noncommunicable diseases and guide treatment decisions to empower disease management in resource-limited settings.

## Acknowledgments

We thank Dr. Kathleen Cormier from the KI histology core for tissue sectioning and staining; Dr. Virginia Spanoudaki and Sarah Elmiligy from the KI Animal Imaging & Preclinical Testing Core for assistance with IVIS imaging; Richard Cook, Heather Amoroso, and Alla Leshinsky from the KI Biopolymers & Proteomics Core for help with the HPLC and mass spectrometry, Dr. Duanduan Ma from the KI Bioinformatics & Computing Core for assistance with transcriptome data analysis. We would also like to thank Drs. Qian Zhong and Edward Tan for helpful discussion on the manuscript.

## Author contributions

L.H., H.E.F., S.N.B. conceived and designed the research. L.H., R.T.Z., C.N. performed research. L.H., H.E.F., S.N.B. wrote and edited the paper.

## Materials and Methods

### Animal models

All animal studies were approved by the Massachusetts Institute of Technology (MIT) committee on animal care (MIT protocol 0417-025-20 & 0217-014-20). All experiments were conducted in compliance with institutional and national guidelines and supervised by Division of Comparative Medicine (DCM) of MIT staff. Female Balb/c and NCr nude mice were kept under standardized housing conditions. We used a sample size of minimum three mice per group for *in vivo* studies, numbers of animals per group were specified in the figure legends. Littermates of the same sex were randomly assigned to experimental and control groups. Establishment of the transplantation mouse models was described below.

### Cell culture

Mouse cell lines MC26-LucF (carrying firefly luciferase, from Kenneth K. Tanabe Laboratory, Massachusetts General Hospital) was cultured in DMEM (Gibco) medium supplemented with 10% (v/v) fetal bovine serum (FBS)(Gibco), 1% (v/v) penicillin/streptomycin (CellGro) at 37 °C and in 5% CO_2_. Human cell lines PC-3 (ATCC^®^ CRL-1435™) were grown in RPMI1640 (Gibco) supplemented with 10% (v/v) FBS and 1% (v/v) penicillin/streptomycin. RWPE1 cells were cultured in Keratinocyte serum-free medium (Gibco) supplemented with 2.5 μg Human Recombinant EGF (rhEGF) and 25 mg Bovine Pituitary Extract (BPE). All cell lines tested negative for mycoplasma contamination.

### Peptide, oligonucleotides and peptide-oligonucleotide conjugates synthesis and characterization

All peptides were chemically synthesized by CPC Scientific, Inc. All oligonucleotides were synthesized by Integrated DNA Technologies, Inc. (IDT). Peptide-oligonucleotides conjugates were generated by copper-free click chemistry. The conjugates were purified on Agilent 1100 HPLC. Mass analysis of the conjugates was performed on a Bruker model MicroFlex MALDI-TOF (matrix-absorption laser desorption instrument time-of-flight). Sequences of all molecules list in Table S1, S3.

### Cas12a fluorescent cleavage assay

LbCas12a (final concentration 100 nM, New England Biolabs) was incubated with 1x NEB Buffer™ 2.1, crRNA (250 nM, IDT) and complementary DNA activators (4 nM unless specifically described, IDT, in solution or spiked in urine) or urine samples collected from experimental animals at 37 °C for 30 min. Reactions were diluted by a factor of 4 into 1x NEB Buffer ™ 2.1 and ssDNA T_10_ F-Q reporter substrate (30 pmol, IDT) into a reaction volume of 60 μL per well. *LbCas*12a activation was detected at 37 °C every 2 min for 3 hours by measuring fluorescence with plate reader Tecan Infinite Pro M200 (λex: 485 nm and λem: 535 nm). Sequences of all oligonucleotides are listed in Table S1. Fluorescence for background conditions (either no DNA activator input or no crRNA conditions) were run with each assay to generate background fluorescence as negative controls. Cas12a ssDNase activity was calculated from the kinetics curve generated on the plate reader, and reflected by the initial reaction velocity (V_0_), which refers to the slope of the curve at the beginning of a reaction.

### Cas12a cleavage assay with lateral flow readout

Samples were prepared similarly and incubated for 30 min at 37 °C as in Cas12a activation assay described above. Reactions were then diluted by a factor of 4 into 1 x NEB Buffer™ 2.1 and ssDNA T_10_ FAM/Biotin reporter substrate (1 pmol, IDT) into reaction volume of 100 μl. Reactions were allowed to proceed at 37 °C for 1-3 hours unless otherwise indicated, and then 20 μl was added to 80 μl of HybriDetect 1 assay buffer (Milenia). HybriDetect 1 lateral flow strips were dipped into solution and intensity of bands was quantified in ImageJ.

### Characterization of DNA activator concentration or length for Cas12a ssDNase activity

To identify the optimal length for detection with Cas12a, we tested truncated native and modified DNA activator lengths from 15-34 nt and found that in the Cas12a fluorescent cleavage assay described above, Cas12a had a peak sensitivity at a native DNA activator length of 24-mer, in which contains PAM sequence and complementary sequence of crRNA. To further explore the robustness of modified DNA activator *in vivo*, phosphorothioate-modified DNA activators with different lengths were injected at 1 nmol in Balb/c mice, respectively, and urine samples were collected after 1 h of injection. Urine samples were applied as DNA activators in the Cas12a fluorescent cleavage assay, Cas12a ssDNase activity triggered by each DNA activator was normalized to that of the 24-mer modified DNA activator.

### Cloning and expression of recombinant nanobodies

Double-stranded gBlocks® gene fragments encoding nanobody of interest with flanking NcoI and BlpI restriction sites, as listed below, were ordered from Integrated DNA Technologies (IA, USA). The gene fragments were cloned into Novogen pET-28a(+) expression vector at NcoI and BlpI restriction sites and transformed into SHuffle^®^ T7 competent *E. coli*. (New England Biolabs Inc., MA, USA). Bacteria colonies encoding the correct gene inserts were confirmed with Sanger sequencing. For subsequent recombinant protein production, a 500 mL secondary culture of SHuffle^®^ T7 competent *E. coli*. encoding nanobody gene of interest was grown in kanamycin-supplemented LB broth at 37 °C from an overnight 3-mL primary culture until optical density at 600 nm (OD600) reached about 0.6-0.8. Nanobody expression was then induced with an addition of isopropyl β-D-1-thiogalactopyranoside (IPTG) (0.4 mM final concentration). The culture was incubated at 27 °C for 24 h after which bacteria were pelleted and stored at −80 °C. Subsequently, the bacteria pellet was thawed on a water bath at 37 °C and lysed with B-PER™ complete bacteria protein extraction reagent (ThermoFisher Scientific, MA, USA). The released nanobody was purified via standard immobilized metal affinity chromatography (IMAC) with Ni-NTA agarose (Qiagen, MD, USA). The product was confirmed *via* SDS-PAGE analysis.

### Synthesis of DNA-encoded synthetic urine biomarker with a nanobody core

Nanobody (2 mg) was incubated at room temperature overnight in Pierce™ immobilized TCEP disulfide reducing gel (7.5 v/v %) (ThermoFisher Scientific, MA, USA) to selectively reduce C-terminal cysteine following a previously established protocol ^31^. The reduced C-terminal cysteine (1 eq.) was reacted with sulfo DBCO-maleimide crosslinker (4 eq.) (Click Chemistry Tools, AZ, USA) in PBS (pH 6.5, 1 mM EDTA) at room temperature for 6 h after which the excess crosslinker was removed with a disposable PD-10 desalting column (GE Healthcare Bio-Sciences, PA, USA). DBCO-functionalized nanobody was further refined on the fast-protein liquid chromatography (FPLC, GE Healthcare). DNA reporter conjugation was performed by incubating DBCO-functionalized nanobody (1 eq.) with azide-functionalized DNA reporter (1.1 eq.) in PBS (pH 7.4) at room temperature for 24 h. Excess DNA reporter was removed via size exclusion chromatography as described above. The product was confirmed via SDS-PAGE analysis and quantified with Quant-iT™ OliGreen™ ssDNA Assay Kit.

### Synthesis of DNA-encoded synthetic urine biomarkers with polymeric cores

Multivalent PEG (40 kDa, eight-arm) containing maleimide-reactive handles (JenKem Technology) was dissolved in 100 mM phosphate buffer (pH 7.0) and filtered (pore size: 0.2 μm). After filtration, the cysteine terminated peptide-DNA conjugates were added at 2-fold molar excess to the PEG and reacted for at least 4 h at room temperature. Unconjugated molecules were separated using size exclusion chromatohraphy with Superdex 200 Increase 10/300 GL column on ÄKTA fast protein liquid chromatography (FPLC, GE Healthcare). The purified nanosensors were concentrated by spin filters (MWCO = 10 kDa, Millipore). Concentration of the nanosensor was quantified Quant-iT™ OliGreen™ ssDNA Assay Kit (ThermoFisher), fluorescence was read on a Tecan Infinite Pro M200 plate reader Quant-iT Oligreen ssDNA Reagent at λex: 485 nm and λem: 535 nm). Particles were stored at 4 °C in PBS. Dynamic light scattering (Zeta Sizer Nanoseries, Malvern Instruments, Ltd) was used to characterized the hydrodynamic diameter of the nanoparticles.

### Transcriptomic and proteomic analysis

RNA-Seq data of human colon adenocarcinoma (285 tumor samples vs 41 normal tissue samples) were obtained from the TCGA Research Network (http://cancergenome.nih.gov). Differential expression analyses were carried out by DESeq2 1.10.1. Proteomic data on the composition of extracellular matrix in human colon cancers and normal colon tissues were obtained by mass spectrometry analysis of ECM components and available from Matrisome (http://matrisomeproject.mit.edu/).

### Establishment of the animal models and urine collection

Balb/c female mice (6-8 wks of age) were inoculated by intravenous (IV) injection with murine cell lines (100k cells/mouse, MC26-Fluc) expressing firefly luciferase. Tumor progression was monitored weekly using IVIS Imaging Systems (IVIS, PerkinElmer). To establish the prostate cancer xenograft model, NCr nude female mice (4-5 wks of age) were inoculated with human PC-3 cell lines (5 million cells per flank, 2 flanks per mouse). Cells were prepared in 30% Corning™ Matrigel™ Membrane Matrix (Thermo Fisher Scientific) and low-serum media (Opti-MEM^®^, Gibco). Tumors were measured weekly and experiments were conducted once flank tumors reached adequate size, which was approximately 5 mm in length or width (~200 mm^3^) or three weeks after inoculation. Tumor volume was calculated by caliper measurement of the length and width of the flank; volume calculation followed the equation *f*_*x*_=IF (length>width, (width^2*length)/2, (length^2*width)/2). For urine analysis, after injection with synthetic biomarkers, mice were placed into custom housing with a 96-well plate base for urine collection. The bladders were voided to collect between 100-200 μL of urine at 1 h post injection. By the end time point of each study, mice were sacrificed and tumor tissues were collected for further analysis.

### Analysis of urinary DNA barcode activated Cas12a cleavage assay

ssDNAs (1 nmol), 5-plex DNA-barcoded PEG sensors (0.2 nmol each by DNA barcode concentration, 1 nmol by DNA barcode concentration in total), or DNA-barcoded nanobody sensors (1 nmol by DNA barcode concentration) were injected into experimental mice via intravenous injection. Urine samples were collected after 1 h and used as DNA activator in Cas12a fluorescent cleavage assay described above. The initial reaction velocity (V_0_) is determined from the slope of the curve at the beginning of a reaction. Mean normalization was performed on V_0_ values to account for animal-to-animal variation in urine concentration. In the Cas12a cleavage assay with fluorescent reporter, Y axis represents MeanNormV_0 Tumor-bearing animals_/MeanNormV_0 control animals._ Then the same urine sample were utilized to perform the Cas12a cleavage assay with LFA readout. Resulting paper strips were aligned and scanned simultaneously, intensity of control and sample bands were quantified from the scanned images in ImageJ.

### Biodistribution and pharmacokinetics studies

Studies were performed in experimental animals with near-infrared dye labeled agents to minimize interference from autofluorescent background. Balb/c mice were intravenously injected with Cy5-labeled modified or native DNA molecules at 1 nmol per mouse, n=3 per condition. Urine samples from each mouse was collected at 30 min, 1, 2, 3, 4 hours after injection. C-met nanobodies were coupled with Sulfo-Cyanine7 NHS ester (Lumiprobe), reacted overnight, purified by spin filtration and injected into PC-3 tumor-bearing nude mice (1.5 nmol dye eq. of protein) *via* i.v. injection. After 24 hours, mice were euthanized and necropsy was performed to remove the tumors, lungs, heart, kidneys, liver, and spleen. Urine, blood and organs were scanned using IVIS Imaging Systems and Odyssey^®^ CLx (LI-COR). Organ fluorescence was quantified in ImageStudio of Odyssey^®^ CLx. Blood circulatory kinetics were monitored in Balb/c mice by serial blood draws at 10 min, 30 min, 120 min and 180 min after i.v. injection of Cy5-labeled DNA or PEG at 1 nmol dye per mouse. Blood for pharmacokinetics measurements was collected using tail vain bleeds. Blood was diluted in PBS with 5 mM EDTA to prevent clotting, centrifuged for 5 min at 5,000 x g, and fluorescent reporter concentration was quantified in 384-well plates relative to standards (LI-COR Odyssey^®^ CLx).

### Histology, Immunohistochemistry (IHC) and Immunofluorescence (IF) studies

Paraffin-embedded tissues were preserved in 4% paraformaldehyde (PFA) overnight and stored in 70% ethanol prior to embedding into paraffin. Snap-frozen tissues were preserved in 2% PFA for two hours, stored in 30% sucrose overnight and frozen in optimum cutting temperature (OCT) compound at −80°C. Snap-frozen lungs were processed through intratracheal injection of 50: 50 OCT in PBS immediately after animal euthanasia. The lungs were slowly frozen with OCT embedding in isopentane liquid nitrogen bath. Samples were sectioned into 6 μm slices and stained for further analysis. For IHC studies, slides were stained with primary antibodies in accordance with manufacturer instructions, followed by HRP secondaries. For IF studies, after blocking with 5% goat serum, 2% BSA, 0.1% Triton-X 100 in PBS for 1 h, sections were stained with a primary antibody in1% BSA in PBS overnight at 4 °C. AlexaFluor conjugated secondary antibodies were incubated at 1 μg/mL in 1% BSA in PBS for 30 min at RT. Slides were sealed with ProLong Antifade Mountants (Thermo Scientific). Slides were digitized and analyzed using an 3D Histech P250 High Capacity Slide Scanner (Perkin Elmer). Antibodies and dilutions used were listed in Table S4.

### RNA extraction and RT-qPCR

PC-3 and RWPE1 cells were cultured and collected after trypsinization. Tissue samples were collected by necropsy after mice were euthanized and were immediately kept in RNAlater RNA Stabilization Reagent (Qiagen, Inc.). RNA from cell pallets or cryogrounded tissue samples was extracted using RNeasy Mini Kit (Qiagen, Inc.). RNA was reverse transcribed into cDNA using BioRad iScript Reverse Transcription Supermix on a Bio-Rad iCycler. qPCR amplification of the cDNA was measured after mixing with Taqman gene expression probes and Applied Biosystems TaqMan Fast Advanced Master Mix (Thermo Scientific) according to manufactory’s instruction. qPCR was performed on a CFX96 Real Time System C1000 Thermal Cycler from Bio-Rad.

### Recombinant protease substrate cleavage assay

Fluorogenic protease substrates with fluorophore (FAM) and quencher (CPQ2) were synthesized by CPC Scientific Inc. Recombinant proteases were purchased from Enzo Life Sciences and R&D Systems. Assays were performed in the 384-well plate in triplicate in enzyme-specific buffer with peptides (1 μM) and proteases (40 nM) in 30 μL at 37 °C. Fluorescence was measured at Ex/Em 485/535 nm using a Tecan Infinite 200pro microplate reader (Tecan). Signal increase at 60 min was used across conditions. Enzymes and buffer conditions were listed in Table S6.

### Protein extraction and tissue lysate proteolytic cleavage assay

Tissue samples were homogenized in PBS and centrifuged at 4 °C for 5 min at 6,000 x g. Supernatant was further centrifuged at 14,000 x g for 25 min at 4 °C. Protein concentration was measured using ThermoFisher BCA Protein Assay Kit and prepared at 2 mg/mL prior to assay. Assays were performed in the 384-well plate in triplicate in enzyme-specific buffer with peptides (1 μM) and cell lysates (0.33 mg/mL) in 30 μL at 37 °C. Fluorescence was measured at Ex/Em 485/535 nm using a Tecan Infinite 200pro microplate reader (Tecan). Signal increase at 60 min was used across conditions.

### Quantification and statistical analysis

Statistical analyses were conducted in GraphPad Prism (Version 8.4). Data were presented as means with standard error of the mean (SEM). Differences between groups were assessed using parametric and non-parametric group comparisons when appropriate with adjustment for multiple hypothesis testing. Results were tested for statistical significance by Student’s *t*-test (parametric) or Mann-Whitney U test (nonparametric) for two group comparisons and ANOVA for multiple group comparisons. Sample sizes and statistical test are specified in the figure legends.

