## Supplemental figures and tables for "CRISPR-Cas-amplified urine biomarkers for multiplexed and portable cancer diagnostics"

for

**Affiliations:**

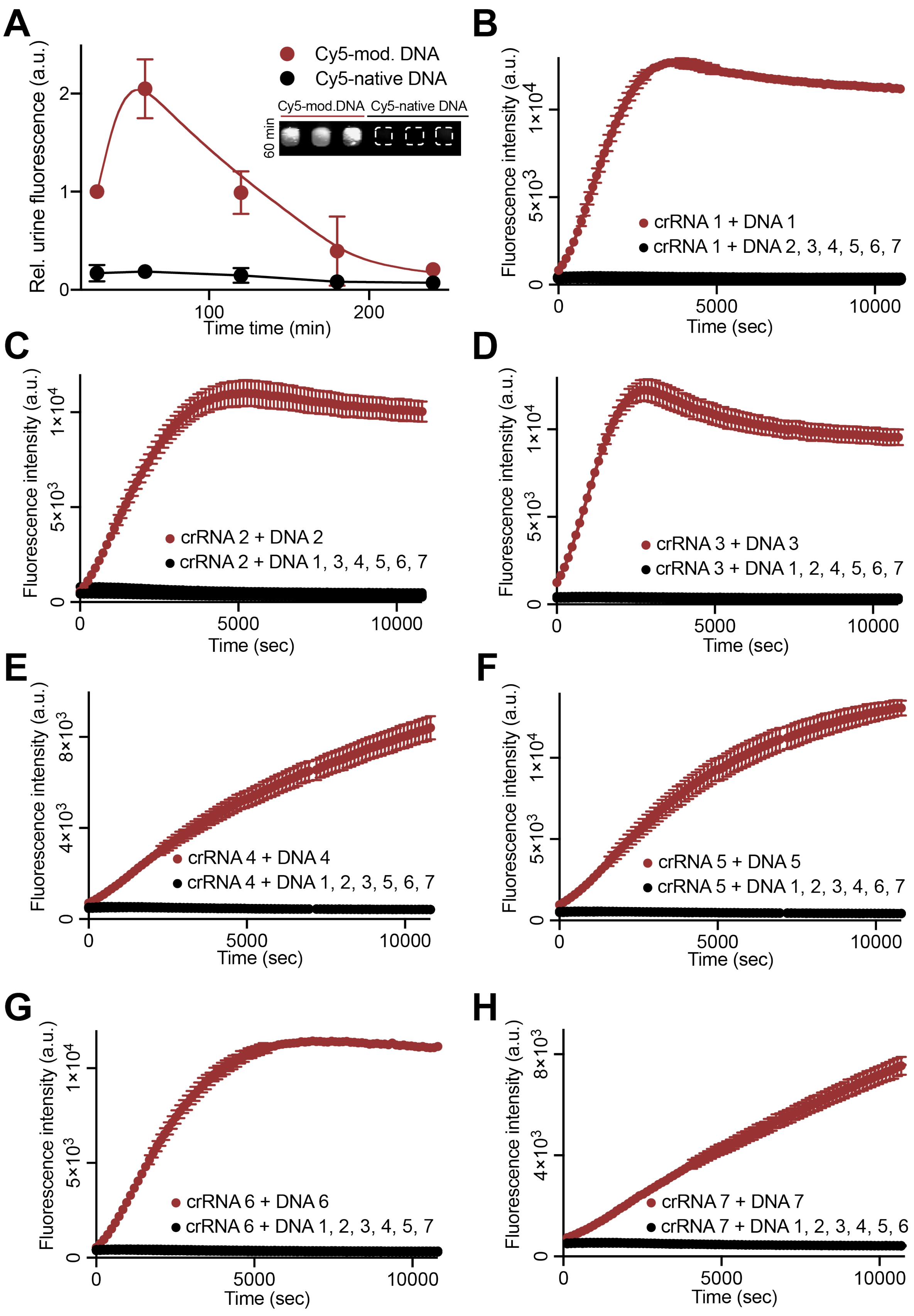
A
Rel. urine fluorescence (a.u.)
Cy5-mod. DNA
Cy5-native DNA
Cy5-mod.DNA
Cy5-native DNA
60 min
Time time (min)
B
crRNA 1 + DNA 1
crRNA 1 + DNA 2, 3, 4, 5, 6, 7
C
crRNA 2 + DNA 2
crRNA 2 + DNA 1, 3, 4, 5, 6, 7
D
crRNA 3 + DNA 3
crRNA 3 + DNA 1, 2, 4, 5, 6, 7
E
crRNA 4 + DNA 4
crRNA 4 + DNA 1, 2, 3, 5, 6, 7
F
crRNA 5 + DNA 5
crRNA 5 + DNA 1, 2, 3, 4, 6, 7
G
crRNA 6 + DNA 6
crRNA 6 + DNA 1, 2, 3, 4, 5, 7
H
crRNA 7 + DNA 7
crRNA 7 + DNA 1, 2, 3, 4, 5, 6
Fluorescence intensity (a.u.)
Time (sec)

**Figure S1. Collateral activity of *Lba*Cas12a activated by different types of DNA activators.** (**A**) The urine signal after systemic administration of modified and native 20-mer DNAs showing amplification kinetics of modified DNA that surpassed the steady-state concentration of its native DNA counterpart. Signal maximized at 1 hour after administration of DNAs. Image shows urine samples on 384-well plate visualized on the LI-COR Odyssey CLx system. Urine fluorescence was normalized to that of the first timepoint of Cy5-modified DNA injected animal (30 min after DNA injection; n=3 per condition). (**B-H**) Trans-cleavage rates of Cas12a upon activation of different modified ssDNA activator-crRNA pairs were determined in the Cas12a fluorescent cleavage assay. Assays were performed with urine samples collected from mice injected with 1 nmol of modified ssDNA activator after 1 h of i.v. administration. The initial reaction velocity ($V_0$) is determined from the slope of the curve at the beginning of a reaction and plotted in Fig. 1D.

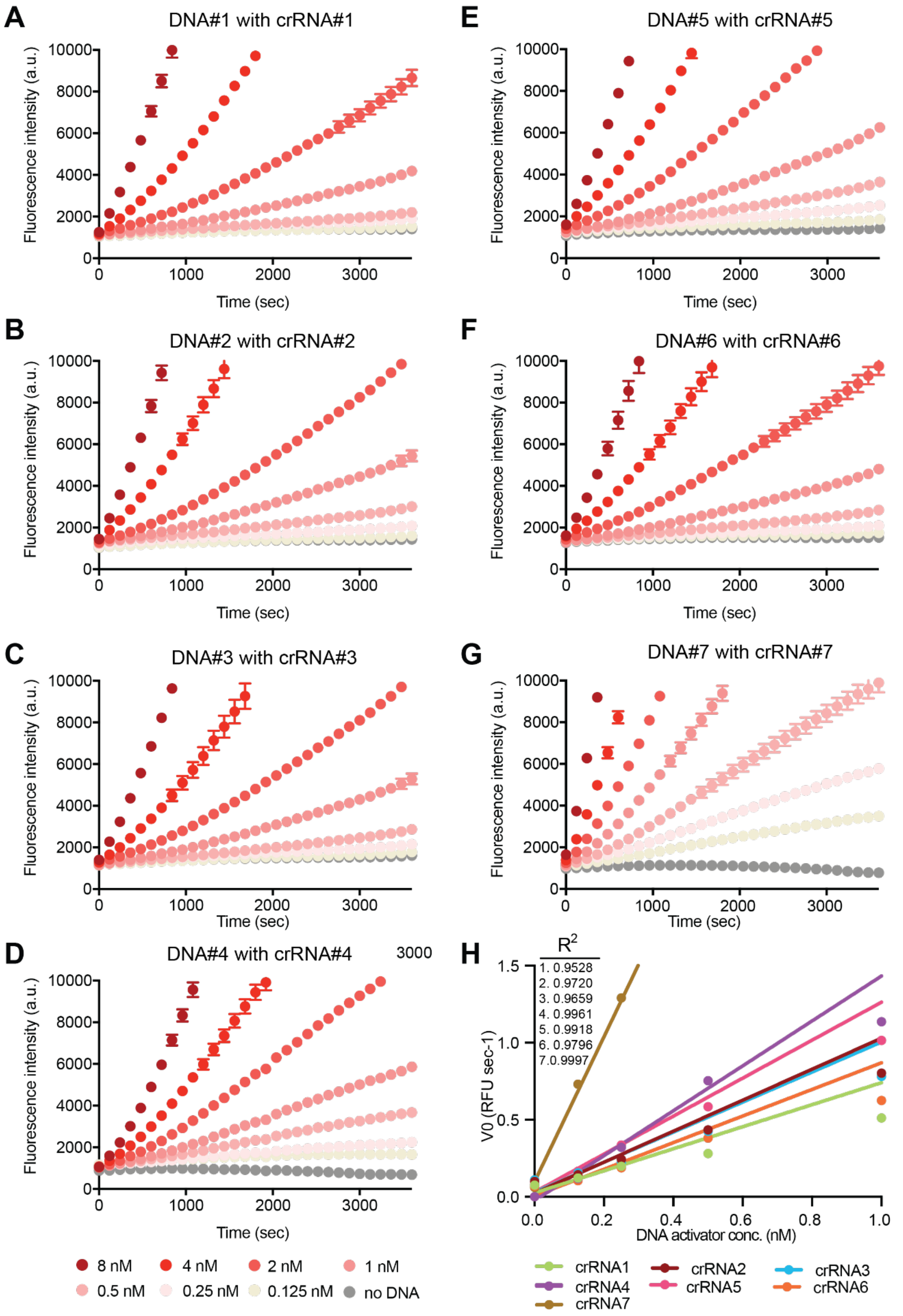

A
DNA#1 with crRNA#1
Fluorescence intensity (a.u.)
Time (sec)
B
DNA#2 with crRNA#2
C
DNA#3 with crRNA#3
D
DNA#4 with crRNA#4
3000
E
DNA#5 with crRNA#5
F
DNA#6 with crRNA#6
G
DNA#7 with crRNA#7
H
R2
1. 0.9528
2. 0.9720
3. 0.9659
4. 0.9961
5. 0.9918
6. 0.9796
7. 0.9997
V0 (RFU sec-1)
DNA activator conc. (nM)
8 nM
4 nM
2 nM
1 nM
0.5 nM
0.25 nM
0.125 nM
no DNA
crRNA1
crRNA2
crRNA3
crRNA4
crRNA5
crRNA6
crRNA7

**Figure S2. Characterizing dose dependence of *Lba*Cas12a activation by DNA activators using fluorescent readout. (A-G**) *Lba*Cas12a catalyzed ssDNA trans-cleavage using phosphorothioate-modified 20- mer ssDNA activators. Trans-cleavage rates of Cas12a upon activation of different modified ssDNA activator-crRNA pairs were determined in the Cas12a fluorescent cleavage assay. Assays were performed with different concentration of modified ssDNA activator (8 nM, 4 nM, 2 nM, 1 nM, 0.5 nM, 0.25 nM, 0.125 nM or 0 nM). (**H**) The initial reaction velocity ($V_0$) is determined from the slope of the curve at the beginning of a reaction in (A-G) and plotted to determine the linear range of assay performance. Linear regions were shown in $V_0$ of reactions for all modified ssDNA activator-crRNA pairs within 1 nM of DNA activators. DNA activator 1, 2, 3, 5, 6 were selected for construction of *in vivo* sensors because of their similarity in assay performance. Sequences of oligonucleotides were shown in Table S1.

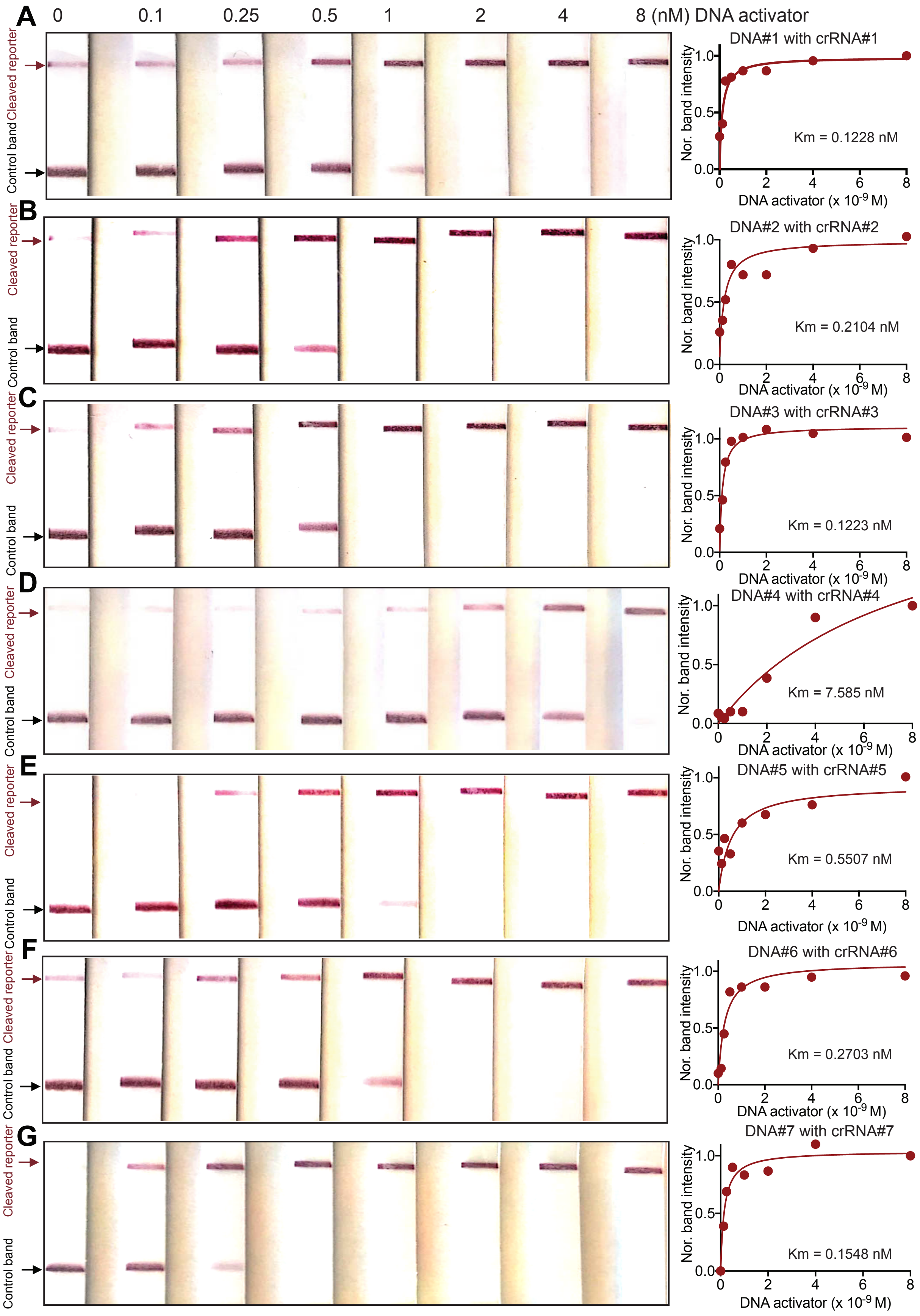
A
0
0.1
0.25
0.5
1
2
4
8 (nM) DNA activator
Cleaved reporter
Control band
DNA#1 with crRNA#1
Km = 0.1228 nM
Nor. band intensity
DNA activator (x 10-9 M)
B
DNA#2 with crRNA#2
Km = 0.2104 nM
C
DNA#3 with crRNA#3
Km = 0.1223 nM
D
DNA#4 with crRNA#4
Km = 7.585 nM
E
DNA#5 with crRNA#5
Km = 0.5507 nM
F
DNA#6 with crRNA#6
Km = 0.2703 nM
G
DNA#7 with crRNA#7
Km = 0.1548 nM

**Figure S3. Characterizing dose dependence of *Lba*Cas12a activation by DNA activators using lateral flow assay.** *Lba*Cas12a catalyzed ssDNA trans cleavage using phosphorothioate-modified 20-mer ssDNA activators. Trans-cleavage rates of Cas12a upon activation of different modified ssDNA activator-crRNA pairs were determined in the Cas12a lateral flow assay. Assays were performed with different concentration of modified ssDNA activator (8 nM, 4 nM, 2 nM, 1 nM, 0.5 nM, 0.25 nM, 0.1 nM or 0 nM) and dual labeled FAM-$T_{10}$-Biotin reporter. Resulting solution was mixed with HybriDetect 1 assay buffer. HybriDetect 1 lateral flow strips were dipped into solution and intensity of cleaved reporter bands was quantified in ImageJ and plotted to fit Michaelis-Menten kinetics. Consistent with the Cas12a fluorescent cleavage assay, linear regions were found within 1 nM of DNA activators for all modified ssDNA activator-crRNA pairs tested. Sequences of oligonucleotides were shown in Table S1.

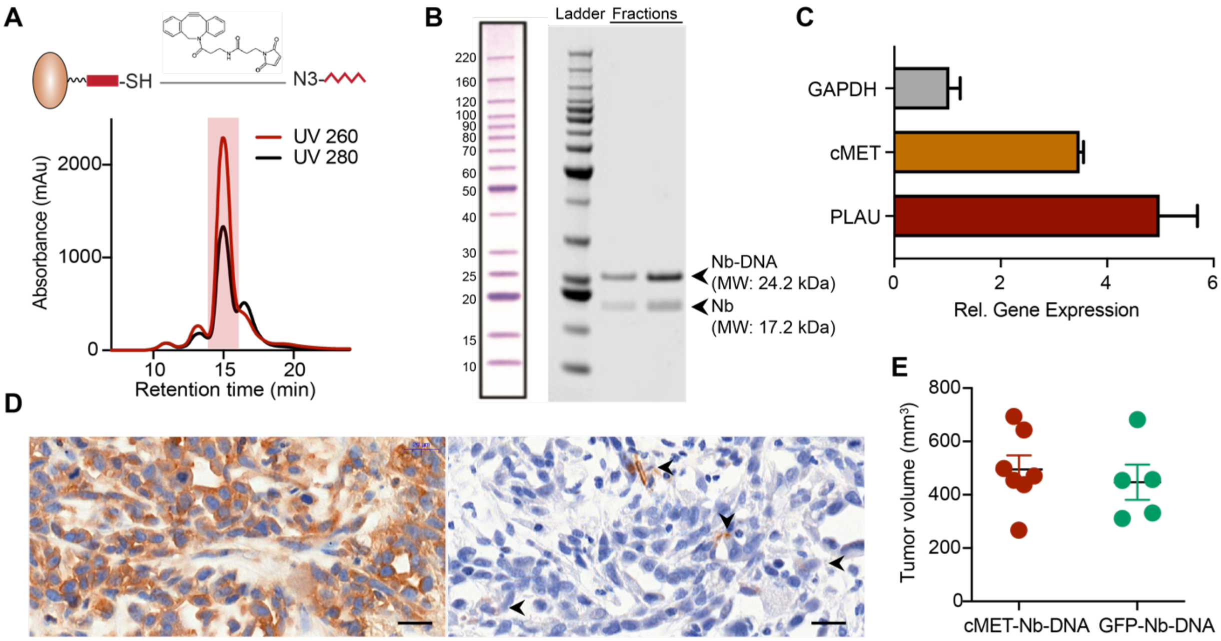


**Figure S4. Characterization of DNA-conjugated nanobody *in vitro* and *in vivo*.** (**A**) Separation of DNA-conjugated cMET nanobody in a size exclusion chromatography. UV 260 nm (red), the elution curve of oligonucleotides; UV 280 nm (red), the elution curve of proteins. Red shed indicates the elution of the DNA-nanobody conjugate that has significant absorbance at both 260 nm and 280 nm. (**B**) SDS-PAGE analysis of the DNA-nanobody conjugate showing predicted molecular weight. (**C**) Increased expression of cMET, the biomarker that the nanobody targets and PLAU, the protease triggers the DNA barcode release, in prostate cancer line PC-3 compared with normal prostate epithelial line RWPE1. (**D**) Immunohistochemical staining of cMET and PLAU in PC-3 flank tumors. Brown, positive staining. Blue, nuclei. Scale bar = 200 µm. (**E**) Caliper quantification of tumor sizes of animals shown in Fig. 3D&E. Tumor-bearing mice were injected with different types of DNA-conjugated nanobodies.

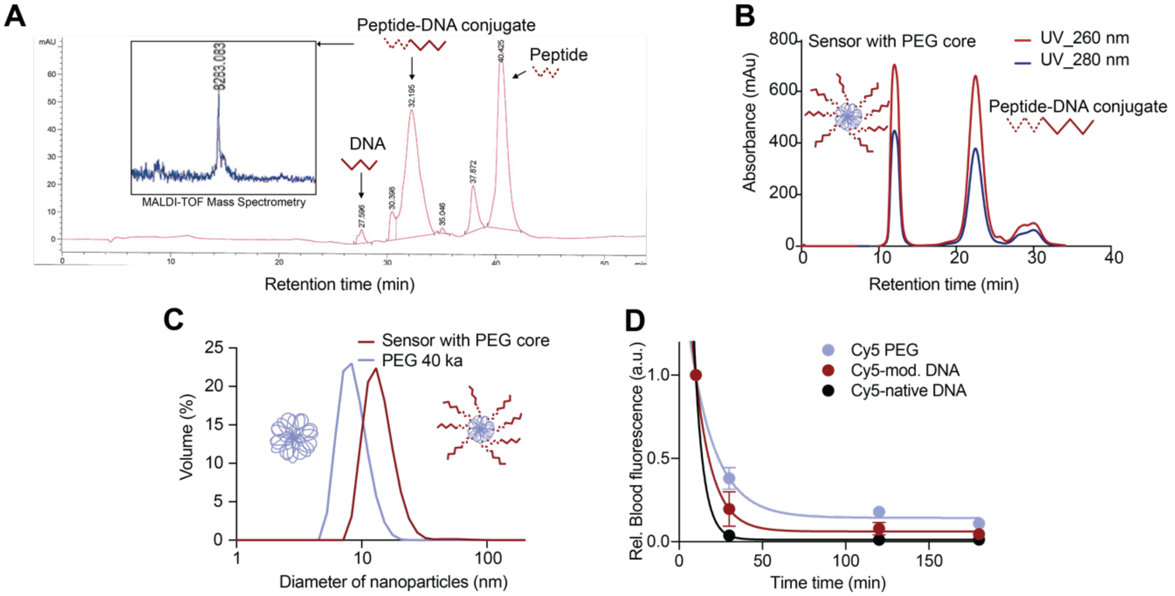


**Figure S5. Characterization of DNA-encoded synthetic urine biomarker built on the polymeric PEG core.** (**A**) Characterization of the representative DNA-PAP7-SUB on a PEG core. HPLC purification of peptide-DNA (PAP7-DNA2) conjugate. The conjugate was analyzed in mass-spectrometry and showed expected molecular weight (8283 Da). (**B**) FPLC purification of sensor showed separation of functionalized sensor and unbounded peptide-DNA conjugate. (**C**) Dynamic light scattering analysis showed increase of particle size from 8.3 nm (PEG core only) and 13 nm (functionalized sensor). (**D**) Plasma half-life shows rapid clearance of native DNA molecules and prolonged half-life of the modified DNA and PEG scaffold in healthy Balb/c mice.

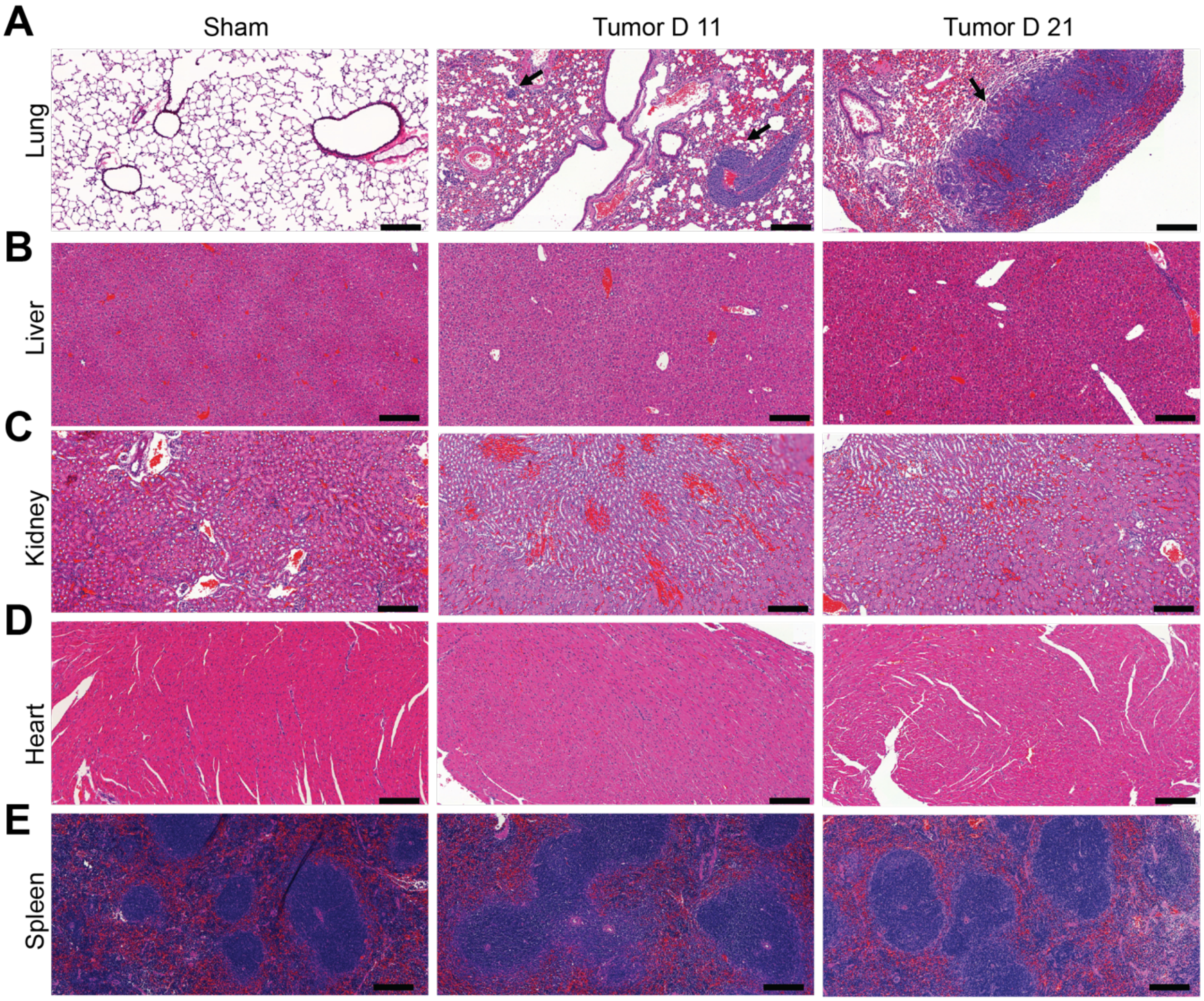


**Figure S6. Histology of major organs of CRC lung metastasis model.** Immunocompetent Balb/c mice were injected with MC26-Fluc cells (tumor) or saline (sham) intravenously. (**A-E**) Organs (lung, liver, kidney, heart and spleen) were collected at 11 and 21 days after administration. Organs were fixed, embedded in paraffin, and stained with hematoxylin & eosin. Study was done with n=3 mice per time point and images from a representative animal are shown. Scale bar = 100 μm. Arrows, tumor nodules in the lung.

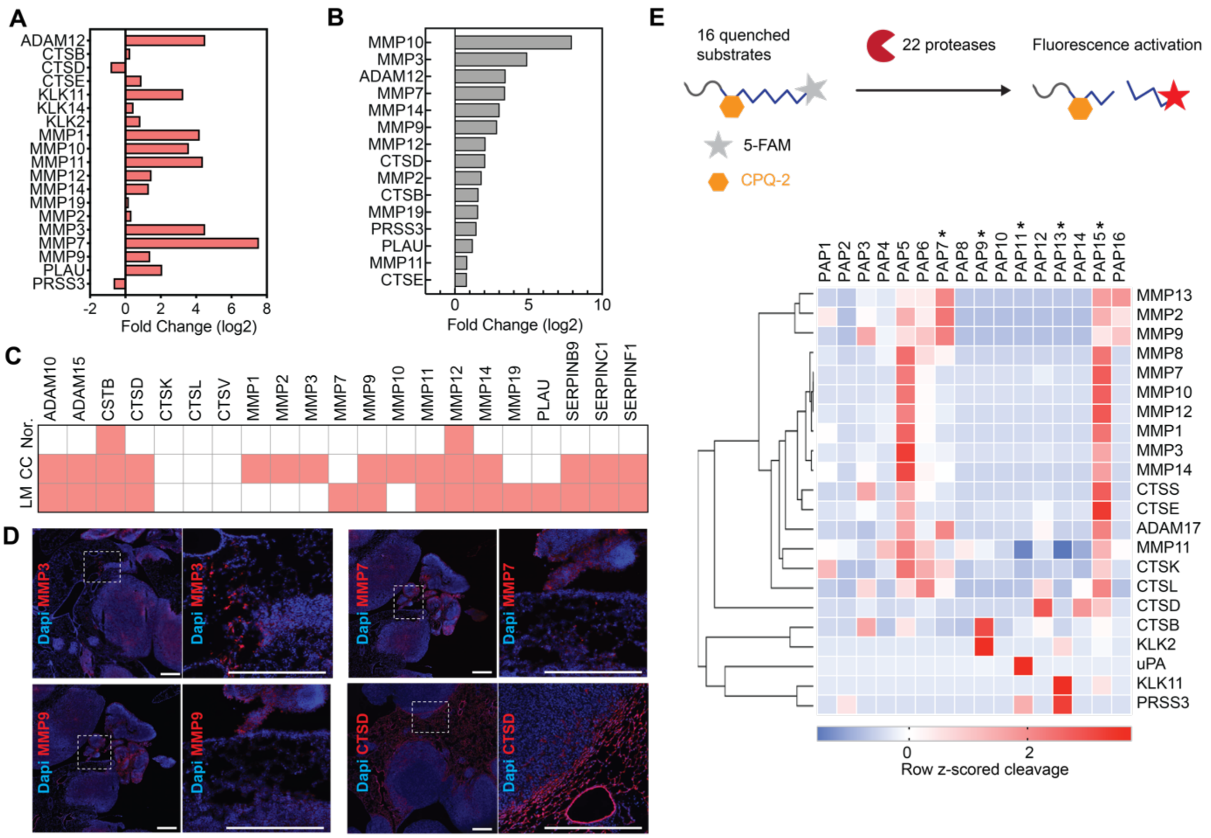


**Figure S7. Identification of deregulated proteases in CRC to select peptide substrates for *in vivo* sensors.** (**A**) Analysis of differentially expressed proteases in CRC samples and normal adjacent tissues. Data available from TCGA. (**B**) RT-qPCR validation of proteases in the tumor-bearing lung from Balb/c mice injected with MC26-Fluc cells in comparison of normal lung from Balb/c mice injected with saline. (**C**) Typical proteases identified in the matrix of primary human colon cancer (CC) and their liver metastases (LM), in comparison of normal colon (Nor.) tissue. Pink, presence; white, absence. Data available from Matrisome project (http://matrisomeproject.mit.edu/). (**D**) Immunofluorescence staining of proteases in the tumor bearing lung tissue sections. Staining of MMP3, MMP7, MMP9 and CTSD is shown in red. Nuclei are counterstained blue with DAPI. Scale bar = 100 μm. (**E**) 16 FRET-paired protease substrates, each consisting of a peptide sequence flanked by a FAM fluorophore and a CPQ-2 quencher, were screened against 22 recombinant proteolytic enzymes. Lower, FRET signal was monitored by kinetic plate reader and the z-scored cleavage rate were subjected to heatmap and Hierarchical Clustering on Morpheus (https://software.broadinstitute.org/morpheus). Asterisk, peptide substrates selected to build *in vivo* sensors because of their broad coverage of metallo, serine and aspartic protease activities.

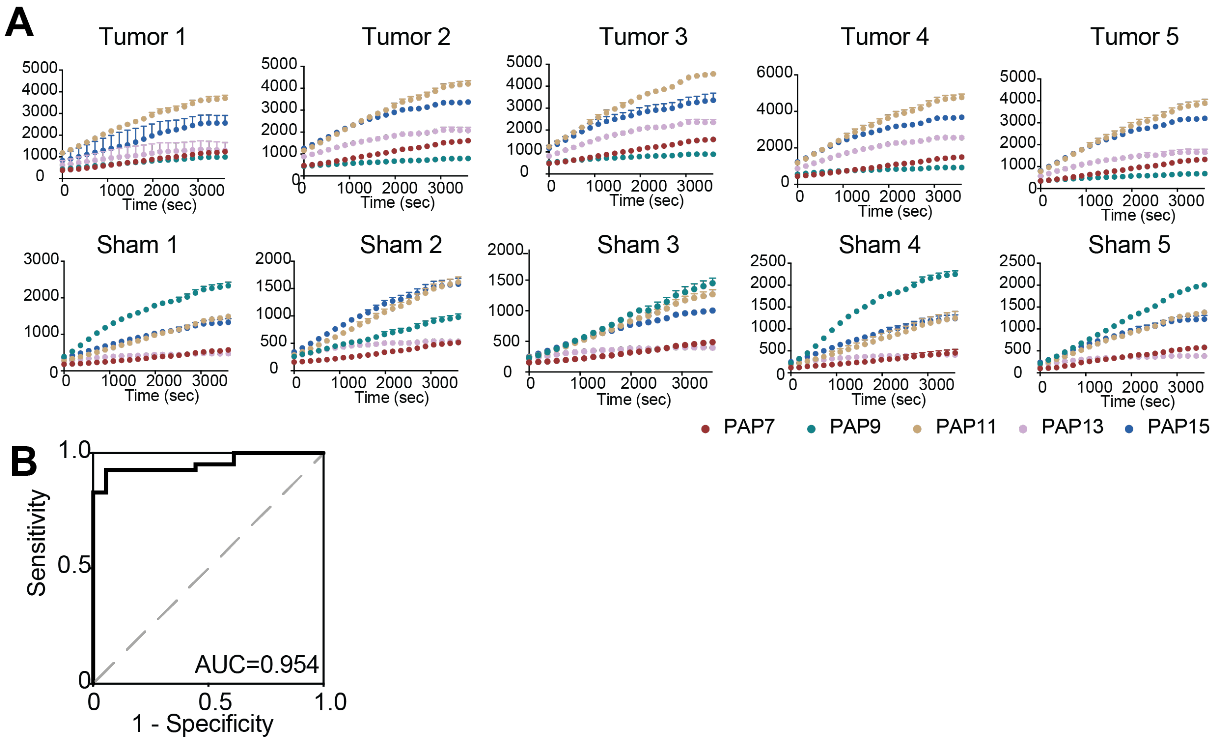


**Figure S8. Classification power of selected peptide substrates in differentiating tumor from normal tissue.** (**A**) Selected FRET-paired protease substrates (PAP 7, PAP 9, PAP 11, PAP 13, PAP 15) were incubated against tissue lysates from tumor bearing lung (tumor, upper) or normal lung (sham, lower) of Balb/c mice (n=5). (**B**) Receiver operating characteristic (ROC) curve analysis of the classification power of the 5 substrates to differentiate tumor vs normal tissue. The area under the curve (AUC) was calculated. Dashed line represents an AUC of 0.5, and a perfect AUC is 1.0.

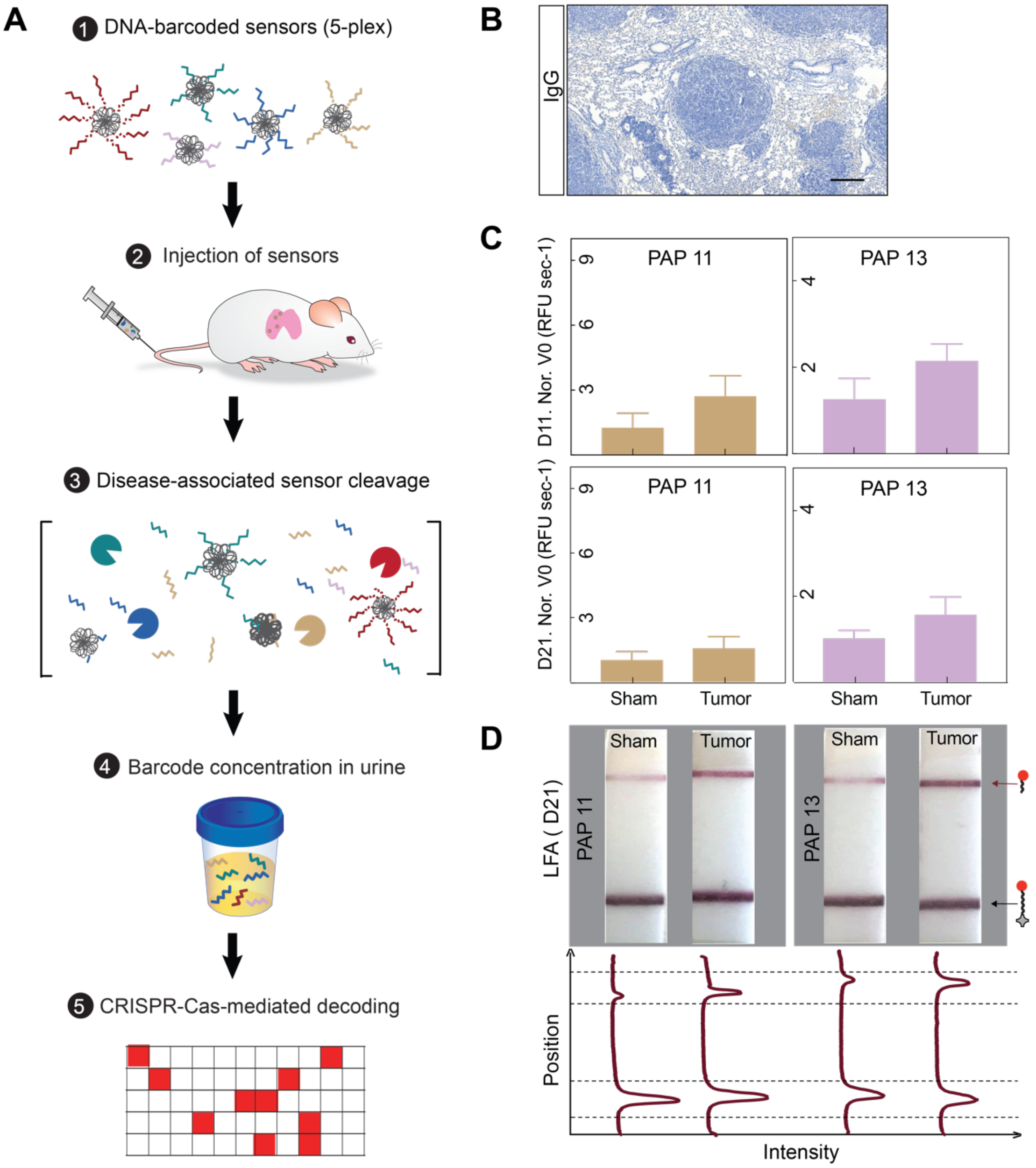


**Figure S9. Multiplexed DNA-encoded synthetic urine biomarkers for disease monitoring.** (**A**) Scheme of the work flow for longitudinal disease monitoring with the multiplexed DNA-encoded synthetic urine biomarkers. (**B**) Immunohistochemistry staining of lung sections of Balb/c mice bearing CRC lung tumor nodules with epitope control antibody. Scale bar = 200 µm. (**C**) 5-plex DNA-SUBs were pooled and administered to Balb/c mice bearing CRC lung tumor nodules and control animals at day 11 or 21 after tumor initiation. All urine samples were collected at 1 h after sensor administration. Two sensors (DNA-PAP11-SUB, DNA-PAP13-SUB) showed an increase in the sets of tumor-bearing mice generated urine signals that were elevated relative to control animals. (**D**) Representative paper strips of the paper-based LFA of Cas12a activated by mouse urine samples collected in (C). Band intensities were quantified using ImageJ. The top peak of the curve shows the presence of the “cleaved reporter” and the bottom peak shows the presence of the “control band”.

**Table S1. Nucleic acid sequences used in this study**

| Name of oligo | Sequence (5'→ 3') |
|---|---|
| crRNA 1 | UAAUUUCUACUAAGUGUAGAUCGUCGCCGUCCAGCUCGACC |
| crRNA 2 | UAAUUUCUACUAAGUGUAGAUGAUCGUUACGCUAACUAUGA |
| crRNA 3 | UAAUUUCUACUAAGUGUAGAUCCUGGGUGUUCCACAGCUGA |
| crRNA 5 | UAAUUUCUACUAAGUGUAGAUCTGTGTTTATCCGCTCACAA |
| crRNA 6 | UAAUUUCUACUAAGUGUAGAUUGAAGUAGAUAUGGCAGCAC |
| crRNA 7 | UAAUUUCUACUAAGUGUAGAUACAAUAUGUGCUUCUACACA |
| ssDNA | TAGCATTCCACAGACAGCCCTCATAGTTAGCGTAACGATCTAAAGTTTTGTCGTC |
| Mod ssDNA | T*A*G*C*A*T*T*C*C*A*C*A*G*A*C*A*G*C*C*C*T*C*A*T*A*G*T*T*A*G*C*G*T*A*A*C*G*A*T*C*T*A*A*A*G*T*T*T*T*G*T*C*G*T*C |
| dsDNA-strand 1 | TAGCATTCCACAGACAGCCCTCATAGTTAGCGTAACGATCTAAAGTTTTGTCGTC |
| dsDNA-strand 2 (complementary) | GACGACAAAACTTTAGATCGTTACGCTAACTATGAGGGCTGTCTGTGGAATGCTA |
| Mod DNA 1 | G*G*T*C*G*A*G*C*T*G*G*A*C*G*G*C*G*A*C*G |
| Mod DNA 2 | T*C*A*T*A*G*T*T*A*G*C*G*T*A*A*C*G*A*T*C |
| Mod DNA 3 | T*C*A*G*C*T*G*T*G*G*A*A*C*A*C*C*C*A*G*G |
| Mod DNA 4 | G*A*G*T*A*A*C*A*G*A*C*A*T*G*G*A*C*C*A*T*C*A*G |
| Mod DNA 5 | T*T*G*T*G*A*G*C*G*G*A*T*A*A*A*C*A*C*A*G |
| Mod DNA 6 | G*T*G*C*T*G*C*C*A*T*A*T*C*T*A*C*T*T*C*A |
| Mod DNA 7 | T*G*T*G*T*A*G*A*A*G*C*A*C*A*T*A*T*T*G*T |
| Dye- DNA 2 | /Cy5/TCATAGTTAGCGTAACGATC |
| Dye- Mod DNA 2 | /Cy5/T*C*A*T*A*G*T*T*A*G*C*G*T*A*A*C*G*A*T*C |
| 10 mer | CGTAACGATC |
| 15 mer | GTTAGCGTAACGATC |
| 20 mer | TCATAGTTAGCGTAACGATC |
| 24 mer | TCATAGTTAGCGTAACGATCTAAA |
| 26 mer | CAGCCCTCATAGTTAGCGTAACGATC |
| 30 mer | CAGCCCTCATAGTTAGCGTAACGATCTAAA |
| 34 mer | GACAGCCCTCATAGTTAGCGTAACGATCTAAAGT |
| Mod 10 mer | C*G*T*A*A*C*G*A*T*C |
| Mod 15 mer | G*T*T*A*G*C*G*T*A*A*C*G*A*T*C |
| Mod 20 mer | T*C*A*T*A*G*T*T*A*G*C*G*T*A*A*C*G*A*T*C |
| Mod 24 mer | T*C*A*T*A*G*T*T*A*G*C*G*T*A*A*C*G*A*T*C*T*A*A*A |
| Mod 26 mer | C*A*G*C*C*C*T*C*A*T*A*G*T*T*A*G*C*G*T*A*A*C*G*A*T*C |
| Mod 30 mer | C*A*G*C*C*C*T*C*A*T*A*G*T*T*A*G*C*G*T*A*A*C*G*A*T*C*T*A*A*A |
| Mod 34 mer | G*A*C*A*G*C*C*C*T*C*A*T*A*G*T*T*A*G*C*G*T*A*A*C*G*A*T*C*T*A*A*A*G*T |
| Mod DNA 1 3'-DBCO | G*G*T*C*G*A*G*C*T*G*G*A*C*G*G*C*G*A*C*G\DBCO |
| Mod DNA 2 3'-DBCO | T*C*A*T*A*G*T*T*A*G*C*G*T*A*A*C*G*A*T*C\DBCO |

| | |
|---|---|
| Mod DNA 3 3’-DBCO | T*C*A*G*C*T*G*T*G*G*A*A*C*A*C*C*C*A*G*G\DBCO |
| Mod DNA 4 3’-DBCO | G*A*G*T*A*A*C*A*G*A*C*A*T*G*G*A*C*C*A*T*C*A*G\DBCO |
| Mod DNA 5 3’-DBCO | T*T*G*T*G*A*G*C*G*G*A*T*A*A*A*C*A*C*A*G\DBCO |
| Mod DNA 6 3’-DBCO | G*T*G*C*T*G*C*C*A*T*A*T*C*T*A*C*T*T*C*A\DBCO |
| Mod DNA 7 3’-DBCO | T*G*T*G*T*A*G*A*A*G*C*A*C*A*T*A*T*T*G*T\DBCO |
| Mod DNA 2 3’- Azide | T*C*A*T*A*G*T*T*A*G*C*G*T*A*A*C*G*A*T*C\Azide |

*, phosphorothioate modification
DBCO, Dibenzocyclooctyne
Cy5, Cyanine 5 dye

**Table S2. Activation of Cas12a with native and modified DNA oligos *in vitro* and *in vivo***
Activation of Cas12a with native and modified DNA oligos were quantified in the Cas12a fluorescent cleavage assay. For 'DNA *in vitro*', 4 nM of DNA activators with different length were added in each reaction. For 'DNA *in vivo*', 1 nmol of DNA activators, native or modified, with different length were injected into healthy Balb/c mice and urine samples collected after 1 h of injection were added in each reaction.

*The initial reaction velocity ($V_0$) refers to the slope of the curve at the beginning of a reaction.

| *Oligo* | Modified DNA *in vitro* ($V_0$)* | Modified DNA *in vivo* ($V_0$) | Native DNA *in vitro* ($V_0$) | Native DNA *in vivo* ($V_0$) |
|---|---|---|---|---|
| *10 mer* | 0.01 | 0.01 | 0.00 | 0.00 |
| *15 mer* | 0.01 | 0.01 | 0.00 | 0.00 |
| *20 mer* | 6.29 | 1.82 | 7.67 | 0.02 |
| *24 mer* | 8.91 | 0.92 | 10.94 | 0.00 |
| *26 mer* | 5.25 | 0.60 | 6.09 | 0.01 |
| *30 mer* | 4.65 | 0.34 | 4.98 | 0.00 |
| *34 mer* | 4.55 | 0.38 | 3.32 | 0.01 |

**Table S3. Peptide and protein sequences used in this study**

| Name of peptide | Sequence (N→ C) |
|---|---|
| Q7-click | K(N3)-ANP-GGPLGVRGKGGC |
| Q9-click | K(N3)-ANP-GG-DPhe-PRSGGC |
| PQ2-click | K(N3)-ANP-GGGSGRSANAKGGC |
| PQ12-click | K(N3)-ANP-GGVPRGGC |
| PQ19-click | K(N3)-ANP-GPVPLSLVMGGC |
| FRET-PAP1 | 5FAM-GGPQGIWGQK(CPQ2)-PEG2-GC |
| FRET-PAP2 | 5FAM-GGLVPRGSGK(CPQ2)-PEG2-GC |
| FRET-PAP3 | 5FAM-GGPVGLIGK(CPQ2)-PEG2-GC |
| FRET-PAP4 | 5FAM-GGPWGIWGQGK(CPQ2)-PEG2-GC |
| FRET-PAP5 | 5FAM-GGPVPLSLVMK(CPQ2)-PEG2-GC |
| FRET-PAP6 | 5FAM-GGPLGLRSWK(CPQ2)-PEG2-GC |
| FRET-PAP7 | 5FAM-GGPLGVRGKK(CPQ2)-PEG2-GC |
| FRET-PAP8 | 5FAM-GGf-Pip-RSGGGK(CPQ2)-PEG2-GC |
| FRET-PAP9 | 5FAM-GGfPRSGGGK(CPQ2)-PEG2-GC |
| FRET-PAP10 | 5FAM-GGf-Pip-KSGGGK(CPQ2)-PEG2-GC |
| FRET-PAP11 | 5FAM-GGGSGRSANAKG-K(CPQ2)-PEG2-GC |
| FRET-PAP12 | 5FAM-GILSRIVGGG-K(CPQ2)-PEG2-GC |
| FRET-PAP13 | 5FAM-GGVPRGG-K(CPQ2)-PEG2-GC |
| FRET-PAP14 | 5FAM-GSGSKIIGGG-K(CPQ2)-PEG2-GC |
| FRET-PAP15 | 5FAM-GPVPLSLVMG-K(CPQ2)-PEG2-GC |
| FRET-PAP16 | 5FAM-GGLGPKGQTGK(CPQ2)-kk-PEG2-C |
| cMET nanobody | MEVQLVESGGGLVQPGGSLRLSCAASGFILDYYAIGWFRQAPGKEREGVLCIDASDDITYYADSVKGRFTISRDNAKNTVYLQMNSLKPEDTGVYYCATPIGLSSSCLLEYDYDYWGQGTLVTVSSGSHHHHHHSPSTPPTPSPSTPP**GSGRSANAK**GGGSC (Clone 4E09, sequence is available from https://patents.google.com/patent/WO2012042026A1/en) |
| GFP nanobody | MAQVQLVESGGRLVQAGDSLRLSCAASGRTFSTSAMAWFRQAPGREREFVAAITWTVGNTILGDSVKGRFTISRDRAKNTVDLQMDNLEPEDTAVYYCSARSRGYVLSVLRSVDSYDYWGQGTQVTVSGSHHHHHHSPSTPPTPSPSTPP**GSGRSANAK**GGGSC (Clone LaG-16)[34] |

Upper case, L-form amino acid;
Lower case, D-form amino acid;
Underlined, rigid linker sequence;
Bolded, PLAU substrate sequence[29];
N3, Azide side chain; ANP, photocleavable linker; 5FAM, N-terminal Fluorescein fluorophore

**Table S4. List of all primary antibodies**

| Antibody | Cat# | Manufacturer | Application | Dilution |
|---|---|---|---|---|
| CTSD | ab75852 | Abcam | IF | 1:100 |
| MMP3 | ab194717 | Abcam | IF | 1:200 |
| MMP7 | ab5706 | Abcam | IF | 1:100 |
| MMP9 | ab38898 | Abcam | IHC, IF | 1:200 |
| PEG | ab190652 | Abcam | IHC | 1:200 |
| PLAU | ab24121 | Abcam | IHC | 1:100 |
| c-Met | ab51067 | Abcam | IHC | 1:100 |
| Cyanine | sc-166895 | Santa Cruz | IF | 1:100 |

**Table S5. List of buffers for proteolytic cleavage assays**

| Enzyme | Manufacturer | Buffer |
|---|---|---|
| MMPs | Enzo | 50 mM TRIS, 10 mM $CaCl_2$, 300 mM NaCl, 20 μM $ZnCl_2$, 0.02% Brij-35, 1% BSA, pH 7.5 |
| ADAMs | Enzo | 10 mM HEPES, 100 mM NaCl, 0.01% Brij-35, 1% BSA, pH 7.4 |
| Cathepsin B | R&D | 25 mM MES, 5 mM DTT, pH 5.0 |
| Cathepsin D | R&D | 0.1 M NaOAc, 0.2 M NaCl, pH 3.5 |
| Cathepsin E | R&D | 0.1 M NaOAc, 0.5 M NaCl, pH 3.5 |
| Cathepsin K | Enzo | 50 mM NaOAc, 1 mM DTT, pH 5.5 |
| Cathepsin L | R&D | 50 mM MES, 5 mM DTT, 1 mM EDTA, 0.005% (w/v) Brij-35, pH 6.0 |
| Cathepsin S | R&D | 50 mM NaOAc, 5 mM DTT, 250 mM NaCl, pH 4.5 |
| uPA/PLAU | R&D | 50 mM Tris, 0.01% Tween 20, 1% BSA, pH 7.4 |
